# PROTAC-Mediated Targeting of IKKβ and NR4A1 for AML Therapy

**DOI:** 10.1101/2025.09.24.678355

**Authors:** Chandra K. Maharjan, Yi Liu, Yufeng Xiao, Bristy R. Podder, Tyler H. Montgomery, Lei Wang, Myung-Chul Kim, Zeng Jin, Seyedehalaleh Anvar, Tanzia Islam Tithi, Vivek M. Shastri, Alexandra M. Stevens, Ryan Kolb, Chen Zhao, Zhijian Qian, Jatinder K. Lamba, Guangrong Zheng, Weizhou Zhang

## Abstract

Acute myeloid leukemia (AML) is an aggressive hematologic malignancy with poor clinical outcomes and limited therapeutic options. Aberrant activation of the IKKβ–NF-κB pathway occurs in approximately 40% of AML cases and contributes to leukemogenesis. However, pharmacological inhibition of IKKβ has been limited by serious toxicities, including neutrophilia. Here we identify IKKβ and NR4A1 as critical drivers of AML progression in certain models and develop a proteolysis-targeting chimera (PROTAC) capable of degrading the proteins. Although NR4A1 has previously been described as a tumor suppressor in AML, our findings demonstrate that NR4A1 exhibits oncogenic functions in some AMLs of the (pro)monocytic lineage. Notably, elevated expression of IKKβ and NR4A1 in AML is associated with poor clinical outcomes, playing non-redundant oncogenic roles in AML. To therapeutically target IKKβ and NR4A1, we designed and synthesized a series of celastrol-based PROTACs that exploit celastrol’s ability to bind both IKKβ and NR4A1. Among these compounds, the lead A9 induces potent cytotoxicity in multiple AML cell lines and primary AML samples through cereblon E3 ligase–dependent degradation of IKKβ and/or NR4A1. In vivo, A9 suppresses leukemia progression in a KMT2A::MLLT3 AML mouse model without inducing neutrophilia, supporting PROTAC-mediated degradation of IKKβ and NR4A1 as a promising therapeutic strategy.

**Plain language summary:** Acute myeloid leukemia (AML) is a predominant form of blood cancer with devastating consequences in both young and adult populations. Despite some advances in AML treatment over the last decade, there is still an unmet medical need to develop new drugs for difficult-to-treat AMLs. We discover that two proteins, namely IKKβ and NR4A1, play important roles in promoting the AML disease and could be associated with more severe AML subtypes. To target and destroy those proteins within AML cells, we employ a novel targeted protein degradation approach, which involved design, synthesis, and screening of PROTAC drugs, to discover a drug possessing pronounced AML cell killing activity by virtue of IKKβ and NR4A1 degradations. The drug also significantly reduces AML progression in mice without any noticeable adverse effects. Our current efforts involve more comprehensively testing the benefit of the drug (and its analogs) in a wide range of clinically relevant AML models.

**Key Points:**

1. IKKβ and NR4A1 are clinically relevant drivers of AML pathogenesis.
2. A novel celastrol-based PROTAC can effectively degrade IKKβ and/or NR4A1 to potentially treat AML.

## Introduction

Significant advances in AML research have facilitated improved diagnosis, cytogenic risk classification, and introduction of new targeted agents for personalized therapies over the past decade [1–3]. The rate of complete disease remission in intermediate- and high-cytogenic risk patients, however, remains low and there is an urgent need for more effective therapies.

NF-κB signaling is constitutively activated in a high percentage of AML patients, promoting cell survival, proliferation, and resistance to therapies by upregulating pro-survival genes and establishing a pro-inflammatory tumor microenvironment (TME) [4–8]. IKKβ is a key kinase for NF-κB activation, whose hyperactivation is a direct consequence of genetic aberrations in AML [5]. IKKβ is an attractive therapeutic target in AML and other cancers, with small molecule inhibitors displaying promising outcomes *in vitro* and *in vivo* [7, 9]; however, systematic IKKβ inhibition raises serious safety concerns, including neutrophilia and systemic inflammation [10, 11].

Proteolysis targeting chimeras (PROTACs) emerge as promising tools for targeted protein degradation [12–14]. PROTAC consists of a protein of interest (POI)-targeting ligand (warhead), an E3 ligase ligand, and a linker. PROTAC brings a POI in proximity of an E3 ligase, facilitating polyubiquitination and proteasomal degradation of the POI. Unlike occupancy-driven small molecule inhibitors, PROTACs act in a catalytic, event-driven manner, which works at much lower (sub-stoichiometric) concentrations and restricts dose-limiting toxicities and potential off-target effects. PROTACs offer tissue selectively since E3 ligases, such as Von Hippel Lindau (VHL) and cereblon (CRBN), have differential expressions across tissues [13, 15].

Celastrol is a natural quinone methide triterpenoid and established IKKβ inhibitor [16–18]. Celastrol-based PROTACs can degrade IKKβ [19]. Celastrol is known to target several other proteins including CHK-1, PI3Kα and NR4A1 [19, 20]. NR4A1 belongs to the steroid-thyroid hormone-retinoid nuclear receptor superfamily and is involved in proliferation, differentiation, apoptosis, metabolism, and development [21, 22]. NR4A1 promotes tumorigenesis through both tumor cell–intrinsic (e.g., cell survival, proliferation, drug resistance, epithelial mesenchymal transition, and metabolic adaptation) and extrinsic (e.g., angiogenesis, inflammation, and T cell dysfunction) mechanisms [20, 23–39]. We demonstrated that a celastrol-based PROTAC suppresses melanoma via NR4A1 degradation [20].

Here, we show that our lead celastrol- and CRBN-based A9 is a robust PROTAC degrader of IKKβ, that is also capable of degrading NR4A1 in AML. Previous studies have demonstrated the tumor suppressive role of NR4A1 in AML [40–42]. Here we demonstrate that NR4A1 and IKKβ have distinct critical roles in promoting AML pathogenesis in several AML cells and can be pharmacologically degraded by A9, which represents a novel therapeutic for AML therapy.

## Materials and Methods (brief summary)

### Animals

7-8 weeks old male C57BL6/J mice were purchased from The Jackson Laboratory. NSG-SGM3 mice were purchased and bred. All animal experiments were conducted in strict compliance with UF Institutional Care and Use Committee (IACUC: 202400000275). The KMT2A::MLLT3 mouse AML model used the GFP-tagged KMT2A::MLLT3 cells from Dr. Chen Zhao at Case Western Reserve University. Detailed description of animal experiments is provided in the supplementary information.

### Cells

MONO-MAC-6, NB-4, MOLM-13, HL-60, and U-937 were shared by Dr Zhijian Qian. HEK293T and A375 cells were acquired from ATCC and cultured under standard conditions. P401 cells (public model name AML001) were from an AML patient (IRB H-3342) characterized by a chromosomal insertion: ins(10;11) [43]. DFAM6855 was purchased from PRoXe (public repository of xenografts) and is a therapy-related AML (IRB 20210216) with a complex karyotype including t(9;11)(KMT2A::MLLT3). Details for drug treatment, CRISPR/Cas9-based knockout of IKKβ and NR4A1, biochemical experiments, and others are included in the supplementary information. All cell lines were summarized (**Supp Table-1**).

### Bioinformatic analysis and Single-cell RNA sequencing analysis

scRNA-seq datasets derived from 7 healthy controls and 10 AML patients (GSE185381) [44] was used and the standard scRNA-seq data processing, quality control, integration, and bioinformatic analysis using R tookit Seurat (v.4.0.6) as previously described [45]. Gene set enrichment analysis (GSEA) was conducted with the escape R package (v.1.6.0) [46]. Correlation analysis was performed using the FeatureScatter function on Seurat. The cor.test() function was used to calculate the statistical significance of the correlation. Library preparation and the bulk RNA sequencing was performed at the University of Florida Interdisciplinary Center for Biotechnology Research (ICBR) Gene Expression and Genotyping Core. Proteomics were performed and analyzed at the IDeA National Resource for Quantitative Proteomics, University of Arkansas for Medical Sciences. Detailed statistics are included under each experiment. Detailed experimental procedures are described in the supplementary information.

## Results

### IKKβ and NR4A1 are bona fide AML targets for therapy

IKKβ is an attractive target for AML therapy [7, 8]. Lasry et al. recently associated proinflammatory signatures in malignant AMLs with poor patient survival [44]. Since IKKβ and NF-κB signaling are central to inflammation [47], we examined *IKBKB* (that encodes IKKβ) expression in healthy bone marrow (BM) cells as well as inflammation high and low BM cells from AML patients in the Single Cell Portal dataset from Lasry et al. [44]. *IKBKB* was strongly expressed in several inflammation-high AML clusters, evidenced by their overlap in UMAP representations (**Fig. 1A**). Correlation analysis between *IKBKB* expression and proinflammatory signature enrichment score in CD34^+^ AML BM cells (in the scRNA-seq dataset by Lasry et al.) revealed a strong positive correlation between the two (**Supp. Fig. 1A–1C**, **Fig 1B**).

**Figure 1:**
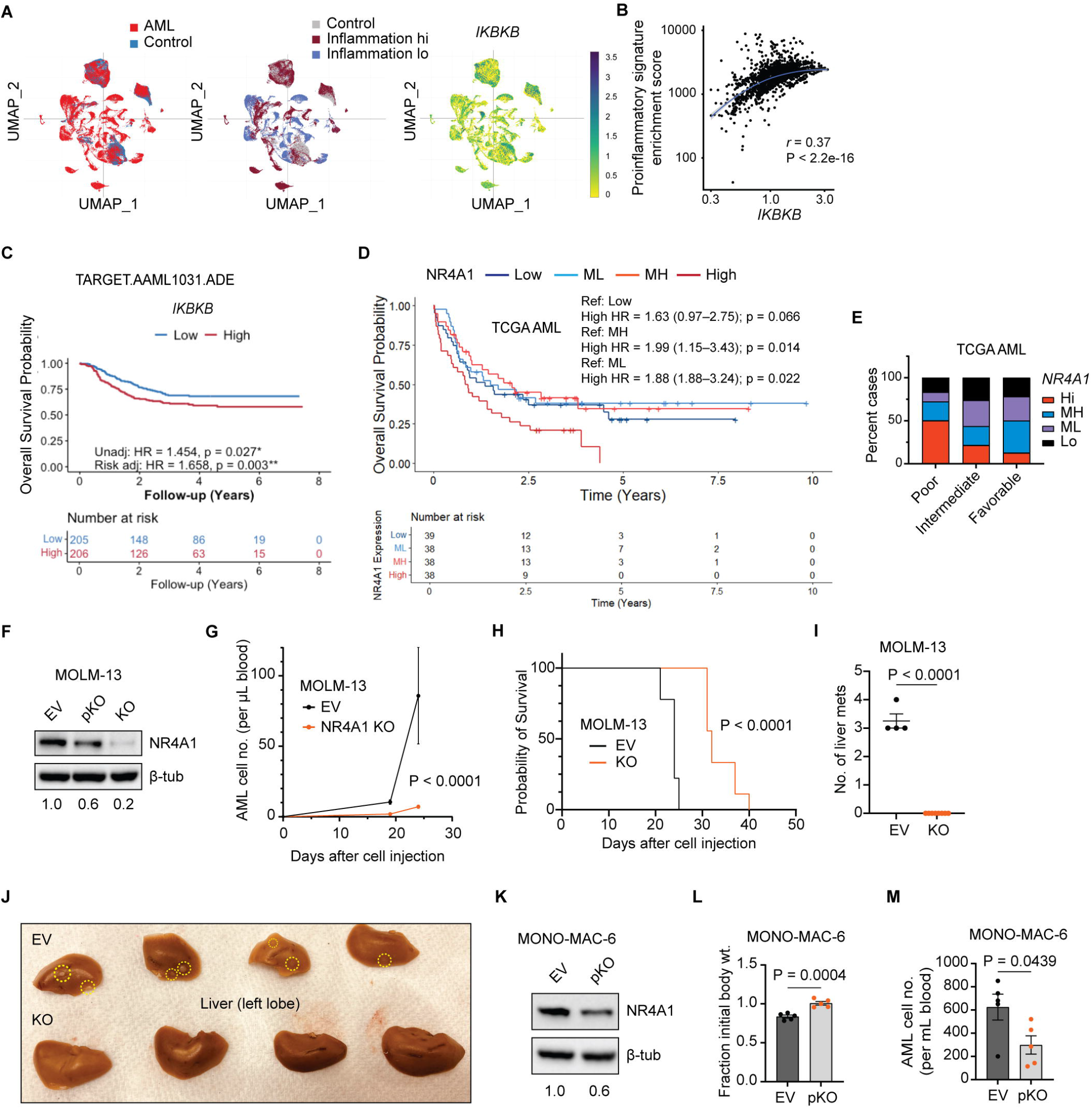
IKKβ and NR4A1 are bona fide AML targets. **(A)** The single cell landscape of adult and pediatric AML derived from a Single Cell Portal analysis of an original dataset generated by Lasry et al., Nature 2022 (PMID 36581735) [44]. Top left—UMAP projection of AML (red) and normal (blue) cells from healthy controls (n=10), adult (n=20) and pediatric (n=22) AML BM cells. Top middle—Depicting healthy control cells and AML cells with inflammation high (hi) and inflammation low (lo) AML cells via UMAP projection. Top right—UMAP projections of *IKBKB* gene expression in healthy control and AML cell clusters. **(B)** Feature scatter plot depicting the positive correlation of *IKBKB* gene expression and enrichment score of proinflammatory signatures in the *CD34*^+^ cells derived from patients with AML. **(C)** Kaplan-Meier plot comparing the overall survival (OS) of *IKBKB* high vs low patients (separated by median) in the ADE (cytarabine, daunorubicin, and etoposide) treatment arm of a patient cohort from the Target- AAML1031 clinical trial. **(D)** TCGA AML patient population divided into four quartiles—high (Hi), moderately high (MH), moderately low (ML), and low—based on NR4A1 expression in AML cells. Kaplan-Meier curves representing overall survival probabilities over time (in days) for each patient quartile in TCGA-AML patient dataset. **(E)** Distribution of AML patients exhibiting varying NR4A1 expressions in each AML cytogenetic risk category presented as stacked bars. **(F)** Pooled *NR4A1* knockout MOLM-13 cells, pKO, were generated via lentiCRISPRv2 technique. Further single clone selection was performed on the pooled cells to obtain the *NR4A1* knockout single clone (KO) as depicted on the western Blotting images. **(G)** 10^6^ EV or *NR4A1* KO MOLM- 13 cells were intravenously injected into the tail vein of NSG-SGM3 mice (n = 9 per group) and their blood were periodically analyzed using flow cytometry to generate the growth curves depicted. **(H)** Kaplan-Meier plot comparing the survival of EV vs *NR4A1* KO MOLM-13 injected mice. **(I)** Number of liver metastases (mets) observed in EV and KO MOLM-13 injected mice. **(J)** Representative images of AML metastases (marked by yellow dotted circles) observed on the mouse livers (left lobes shown); the mice were euthanized at end point defined by low BCS (body weight loss, severe dehydration, and hind limb paralysis). **(K)** Western Blotting images showing NR4A1 protein levels in EV control vs NR4A1 pooled knockout (pKO) MONO-MAC-6 cells generated via lentiCRISPRv2 technique. **(L–M)** 0.5x10^6^ EV or *NR4A1* pKO MONO-MAC-6 cells were injected into the tail vein of NSG-SGM3 mice (n = 7 per condition) and their body weights relative to the initial weights **(L)** and leukemic burden in the blood **(M)** after 34 days of cell injection are shown (n = 5 mice were alive per group). Data from animal experiments are presented as mean +/- SEM. Unpaired two-tailed T-tests were performed for statistical analysis in **I**, **L**, and **M**. Unadjusted and risk group–adjusted hazard ratios were obtained using Cox proportional hazard models in **C**, and **D**. Statistical significance of AML cell growth in vivo was determined by two-way ANOVA (mixed effects analysis) in **G**. Numbers below the Western Blotting indicates relative expression normalized to the control samples. *P < 0.05, **P < 0.01, ***P < 0.001, ****P < 0.0001.

Furthermore, among tested pediatric AML cohorts, higher *IKBKB* expression was associated with poorer overall survival (OS) and event-free survival (EFS) in the standard chemotherapy arm of the Target-AAML1031 trial when adjusted for risk groups (**Fig. 1C, Supp. Fig. 1D–1F**). To determine the function of IKKβ in AML cells, we knocked down *IKBKB* in MONO-MAC-6 cells and found that acute depletion of IKKβ significantly reduced AML cell proliferation (**Supp. Fig. 1G–1I)** without inducing cell death (**Supp. Fig. 1J–1K**).

Considering the oncogenic role of IKKβ in AML, we investigated the therapeutic potential of celastrol-based PROTACs designed to degrade IKKβ. We and others have shown that celastrol-based PROTACs degrade NR4A1, a known celastrol target [19, 20]. BM cells of AML patients exhibit varying levels of NR4A1 (**Supp. Fig. 2A**), but there was no correlation with proinflammatory signature score (**Supp. Fig. 2B**) [44]. Overall survival analysis of patients in the TCGA-AML dataset demonstrated that patients with higher NR4A1 expression associated with a trend towards decreased survival probability compared to those with low expression, without being significantly different (**Supp. Fig. 2C**). Upon further subdivision into quartiles (Q1–Q4) based on NR4A1 expression, those with highest expression quartile had the shortest survival followed by low, moderately high (MH), and moderately low (ML) (**Fig. 1D**). Highest NR4A1 expression quartile was significantly associated with poor outcome compared to MH and ML groups but not with the lowest expression quartile (high HR = 1.63, 95% CI [0.97-2.75]; p = 0.066). Both MH and ML groups had similar survival to those in the low expression group (MH HR = 0.84, 95% CI [0.47-1.48]; p = 0.54 and ML HR = 0.85, 95% CI [0.48-1.49]; p = 0.56). While the difference between survival in high and low expression groups did not reach conventional statistical significance either, the trend of poorest survival in the high expression group remained the same as in the median cut-off analysis. Furthermore, NR4A1 expression, when used as a continuous variable, was significantly associated with poor overall survival (HR = 1.14, 95% CI [1-1.3], p = 0.0493).

**Figure 2:**
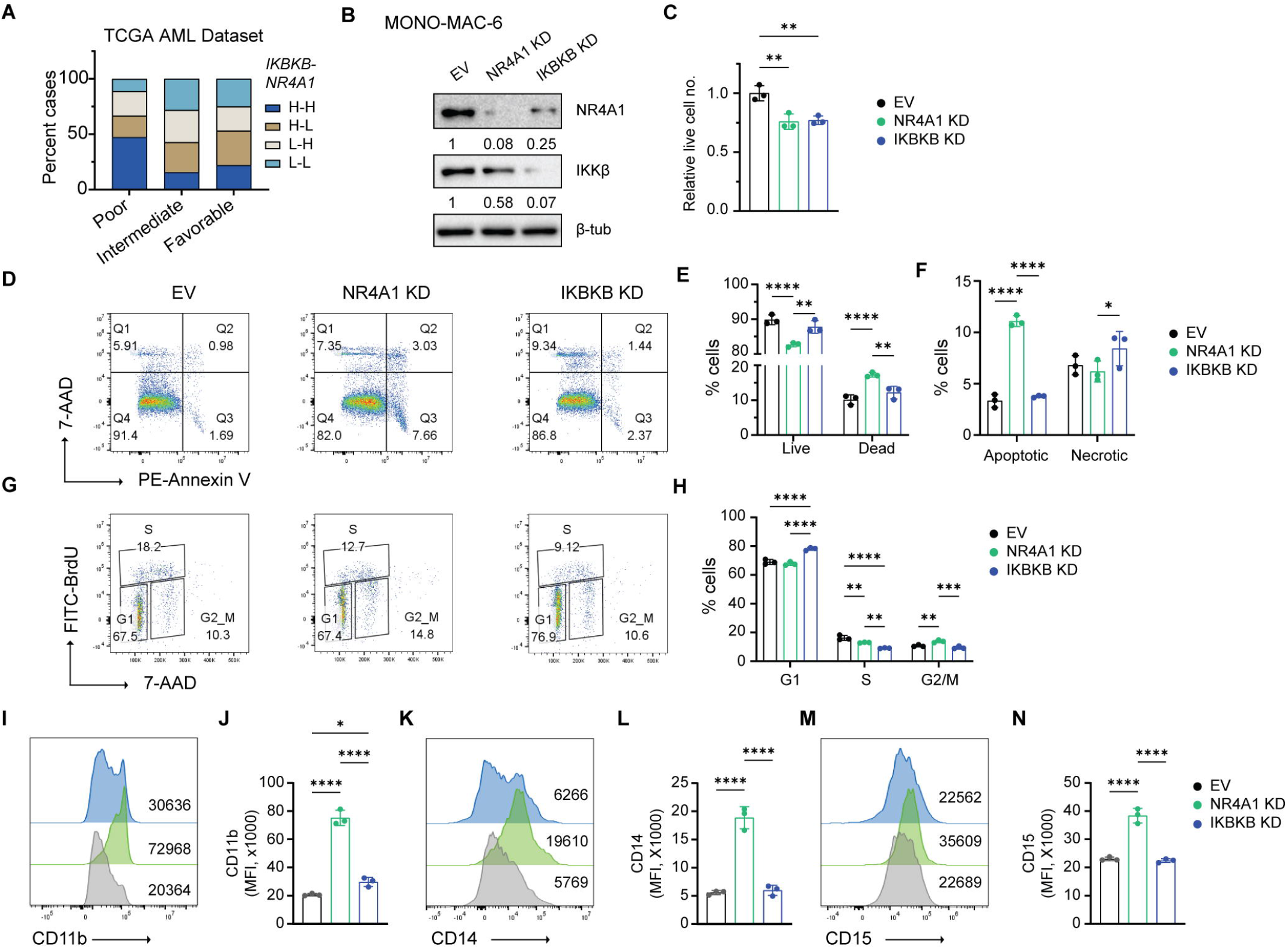
NR4A1 and IKKβ positively regulate each other and play distinct oncogenic roles in certain AML models. **(A)** AML patients in the TCGA database were grouped based on their expression of *NR4A1* and *IKBKB*—both high (H-H), high IKBKB and low NR4A1 (H-L), low IKBKB and high NR4A1 (L-H), both low (L-L). Distribution of AML patients from these 4 subgroups in each AML cytogenetic risk category presented as stacked bars. **(B–M)** EV, *NR4A1* KD, and *IKBKB* KD MONO-MAC-6 cells were generated by transducing corresponding lentiviral shRNAs. **(B)** Western Blotting images of NR4A1 and IKKβ across the above KD conditions in MONO-MAC-6 cells harvested 48 h after terminating lentiviral infection. Numbers indicate the relative protein levels quantified by densitometric analysis in ImageJ. **(C)** Live cell counts of KD cells (on Day 5 post termination of lentiviral infection) relative to EV MONO-MAC-6 cells quantified by trypan blue exclusion method. n = 3 (EV and KD). **(D–F)** Annexin V/7-AAD labeling of EV, *NR4A1* KD, and *IKBKB* KD MONO-MAC-6 cells to quantify the percentages of live (Q4), dead (Q1 + Q2 + Q3), apoptotic (Q2 + Q3), and necrotic (Q1) cells by flow cytometry as shown by representative images in **D**. Cumulative data depicted as bar diagrams in **E** and **F**. **(G–H)** BrdU cell proliferation assay was performed for EV and various KD conditions after an initial 2 h BrdU treatment. Representative images show the distribution of cells at different phases of the cell cycle in **G** and cumulative data summarized in a bar graph in **H**. **(I–N)** Evaluation of mature monocytic differentiation markers, CD11b **(I–J)**, CD14 **(K–L)**, and CD15 **(M–N)**, in EV, *NR4A1* KD, and *IKBKB* KD MONO-MAC-6 cells. Representative histograms with numbers indicating mean fluorescence intensities (MFI) are followed by bar graphs showing cumulative data for n = 3 biological replicates. Data are presented as mean +/- SD. Statistical significance determined using one-way ANOVA in **C**, **J**, **L**, and **N** and two-way ANOVA in **E**, **F**, and **H** followed by Tukey’s multiple comparison tests. *P < 0.05, **P < 0.01, ***P < 0.001, ****P < 0.0001. Only significant comparisons are marked.

The 2022 European Leukemia Network defined risk classification of AML patients based on genetic abnormalities [1, 48]. We found that 50% of AML patients in the poor cytogenetic risk group had high NR4A1 expression, whereas most patients with favorable prognosis had low to moderate NR4A1 expression (**Fig. 1E**). Direct comparison of NR4A1 High vs Low using Hallmark GSEA showed significant enrichment of inflammatory pathways (most prominently TNFα–NF-κB signaling) and several oncogenic pathways in the NR4A1 High group (**Supp. Fig. 2D–2F)**. To test NR4A1’s function in AML, we knocked down *NR4A1* in MOLM-13 human AML cell line (**Supp. Fig. 2G**), noting that we could only achieve marginal but consistently around 20% total protein reduction from pooled culture. We found that acute shRNA-mediated depletion of NR4A1 induces AML cell death (**Supp. Fig. 2H–1I**) as well as inhibits proliferation apparently via cell cycle arrest at G2/M phase (**Supp. Fig. 2J–2K**). Moreover, upregulation of mature monocytic differentiation markers, CD14 and CD15, was observed in *NR4A1* silenced MOLM-13 cells (**Supp. Fig. 2L–2M**). To investigate whether these in vitro results translate to in vivo, we knocked out (KO) *NR4A1* from MOLM-13 cells (**Fig. 1F**). MOLM-13 *NR4A1* KO cells grew slower compared to their EV counterparts (**Supp. Fig. 2N**). Consistently, studies *in vivo* showed slower disease progression (**Fig. 1G**), improved survival (**Fig. 1H**), and complete loss of liver metastasis in NSG-SGM3 mice injected with *NR4A1* KO MOLM-13 cells compared to empty vector (EV) counterparts (**Fig. 1I–1J**). *NR4A1* knockdown (KD) in another MONO-MAC-6 AML cell line led to much more efficient reduction in NR4A1 protein (**Supp. Fig. 2O**), which resulted in pronounced cell death (**Supp. Fig. 2P–2Q**) as well as inhibition of proliferation with both G1 and G2/M arrest (**Supp. Fig. 2R–2S**). In vivo, partial KO of *NR4A1* in MONO-MAC-6 cells (**Fig. 1K**) also led to reduced disease index in xenografted NSG-SGM3 mice, as indicated by improved body weight (**Fig. 1L**) and reduced leukemic burden in the blood (**Fig. 1M**). There was a modest trend towards improved survival in the pKO group, albeit not statistically significant (**Supp. Fig. 2T**). Considering NR4A1 has also been shown to have a tumor suppressive role in certain AML models[40–42], our results point toward a dichotomous NR4A1 role that depends on the disease subtype in AML pathogenesis.

### IKKβ and NR4A1 exhibit mutual positive regulation in certain AML models and play distinct oncogenic roles in AML pathogenesis

We stratified TCGA-AML patients into four groups based on IKKβ and NR4A1 expression in leukemic cells—both high (HH), high IKKβ and low NR4A1 (HL), low IKKβ and high NR4A1 (LH), and both low (LL) —and examined their distribution across cytogenetic risk categories. Notably, patients with high IKKβ and NR4A1 expression constitute nearly half of those with potentially poor outcomes (**Fig. 2A**), suggesting this subset may benefit most from dual targeting of IKKβ and NR4A1. To comparatively evaluate NR4A1 and IKKβ’s oncogenic functions in AML, we knocked down *NR4A1* or *IKBKB* (with EV as a control) in MONO-MAC-6 cells within the same experimental setting. Western Blotting results revealed reciprocal positive regulation between NR4A1 and IKKβ at the protein level (**Fig. 2B**). *NR4A1* KD and *IKBKB* KD both led to a significant reduction in viable AML cell count by day 5 post shRNA transduction (**Fig. 2C**). Functional studies demonstrated that acute loss of NR4A1 primarily induces cell death via apoptosis (**Fig. 2D–2F**); while modestly reducing proliferation (**Fig. 2G–2H**) in MONO- MAC-6 cells, with significant upregulation of mature monocytic differentiation markers, CD11b, CD14 and CD15 (**Fig. 2I–2N**). We also observed that *NR4A1* silencing significantly increases the percentage of CD11b-high and CD14-high MONO-MAC-6 cell populations (**Supp. Fig. 3A–3D**). *IKBKB* KD, on the other hand, strongly inhibited cell proliferation (**Fig. 2G-2H**) with limited effects in cell survival (**Fig. 2D–2F**) and differentiation (**Fig. 2I–2N, Supp. Fig. 3A–3D**). Considering NR4A1 and IKKβ have non-overlapping roles in AML pathogenesis, development of drugs that can target both the proteins holds promise for effective AML therapy. Consistently, knockdown (KD) of *NR4A1*, *IKBKB*, or both genes in NB-4 cells reduced their viability, with the largest effect in the dual KD condition (**Supp. Fig. 3E–3G**). We also observed that NR4A1 positively regulates IKKβ at the protein level; however, the reverse was not true (**Supp. Fig. 3F**), suggesting a cell type dependent regulation of IKKβ to NR4A1 protein.

**Figure 3:**
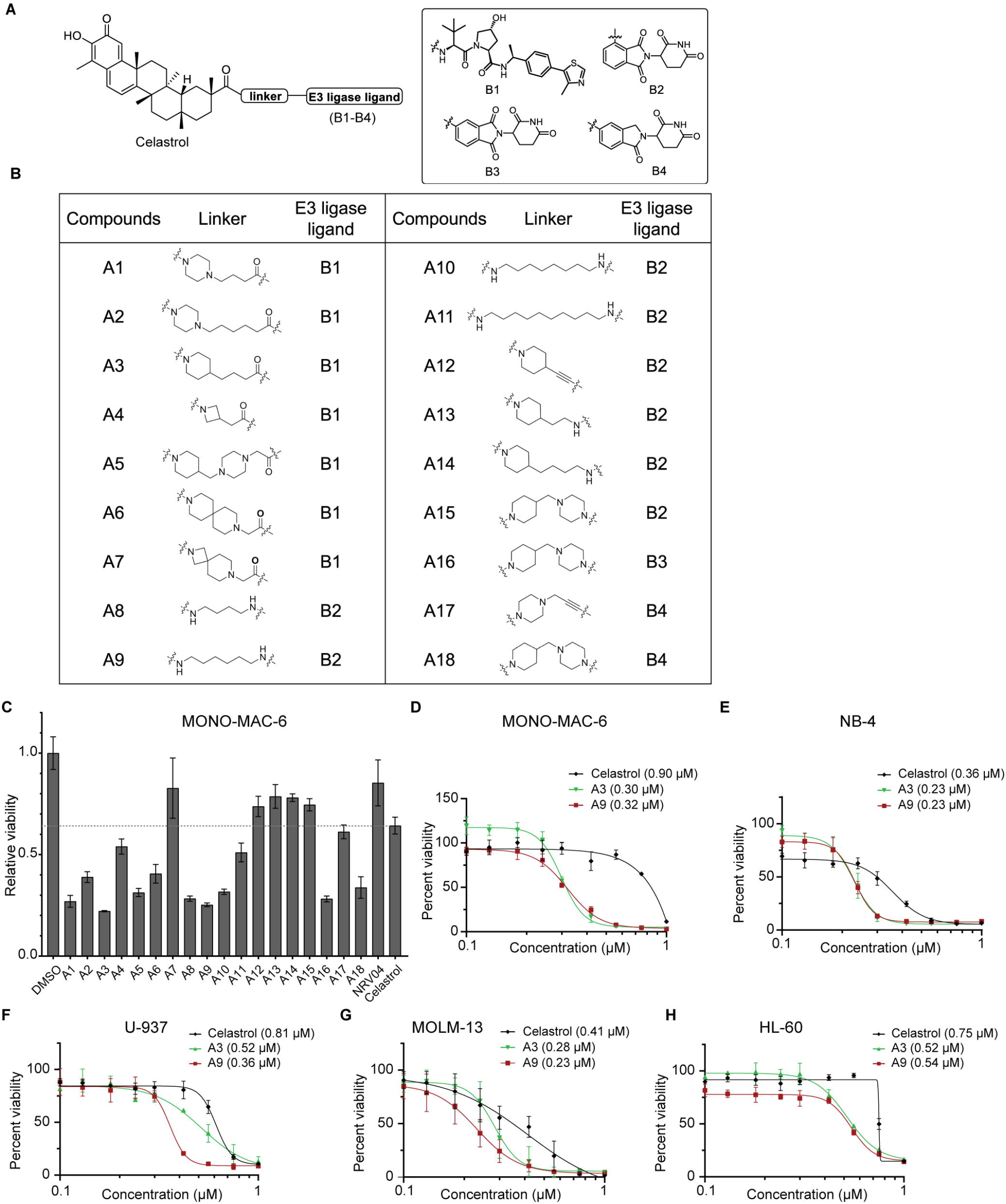
Two celastrol based PROTACs, A3 and A9, were most potent in reducing AML viability. **(A–B)** Panel of celastrol based PROTACs synthesized and screened in AML cells. **(A)** Overall strategy employed in the synthesis of our PROTAC drugs, A1–A18. B1 is a VHL ligand, B2–B4 are CRBN ligands; B2 and B3 represent pomalidomide linked to celastrol through different positions whereas B4 represents lenalidomide. **(B)** Structures of linkers used in A1– A18 PROTAC synthesis. Data are presented as mean +/- SEM. **(C)** A panel of celastrol-based PROTACs were screened for AML cell killing in MONO-MAC-6 cell line via MTS assay. Bars represent viabilities of cells treated with celastrol PROTACs or celastrol for 48 h relative to DMSO. **(D–H)** Comparison of A3 and A9 kill curves with that of celastrol in MONO-MAC-6 **(D)**, NB-4 **(E)**, U-937 **(F)**, MOLM-13 **(G)**, and HL-60 **(H)** AML cells. Data presented as mean +/- SD. IC50 values of drug response curves indicated inside parentheses within the figure legends.

### Two celastrol-based PROTACs, A3 and A9, most strongly reduced AML cell viability

Our group previously identified a potent NR4A1 degrader, NR-V04, by constructing and screening a celastrol-based PROTAC library incorporating E3 ligase ligands and linkers (patent # WO-2022072094-A2) [20]. To discover PROTACs capable of killing AML cells and potentially degrading both NR4A1 and IKKβ, we performed a cell viability-based screening of our PROTAC library (**Fig. 3A–3B**) against two AML cell lines, MONO-MAC-6 (**Fig. 3C**) and NB-4 (**Supp. Fig. 3H**). This library included PROTACs featuring both CRBN- and VHL-ligands, joined by various linkers (**Fig. 3A–3B**). MTS assay results identified A3 and A9 as having the highest AML killing activity across both cell lines (**Fig. 3C, Supp. Fig. 3H**). We then treated MONO-MAC-6, NB-4, U-937, MOLM-13, and HL-60 AML cells with varying concentrations (0.1–1 µM) of A3, A9, and celastrol (**Fig. 3D–3H**). Both A3 and A9 demonstrated greater potency than celastrol in killing AML cells with nanomolar IC50 values. We observed AML cell line-specific differences in the dose response curves, which could be related to the variations in IKKβ and NR4A1 expression levels (**Supp. Fig. 3I)** as well as functional dependencies of the cells on these putative targets.

### A9 induces death and inhibits proliferation in AML cells

Effective E3 ligase binding is critical for PROTAC-mediated protein degradation. Structurally, A9 consists of celastrol linked to a CRBN E3 ligase ligand (pomalidomide) through a six-carbon chain (**Fig. 4A**), while A3 links celastrol to a VHL ligand through a piperidine ring and short hydrocarbon chain (**Supp. Fig. 4A**). To test whether AML cell killing by A9 and A3 requires binding to their corresponding E3 ligases, we synthesized negative controls, A9 NC and A3 NC. The NH group of glutarimide in the CRBN ligand was protected by a methyl group in A9 NC (**Fig. 4A**) whereas a diastereoisomeric OH group was introduced on the hydroxyproline moiety of the VHL ligand in A3 NC (**Supp. Fig. 4A**) to diminish the E3 binding capacity. A9 NC was less potent than A9 in killing NB-4 and MONO-MAC-6 cells (**Fig. 4B**), while the kill curves of A3 and A3 NC overlapped (**Supp. Fig. 4B**). These results suggest that cytotoxic functions of A9, but not A3, in AML is in part dependent on its E3 ligase binding.

**Figure 4:**
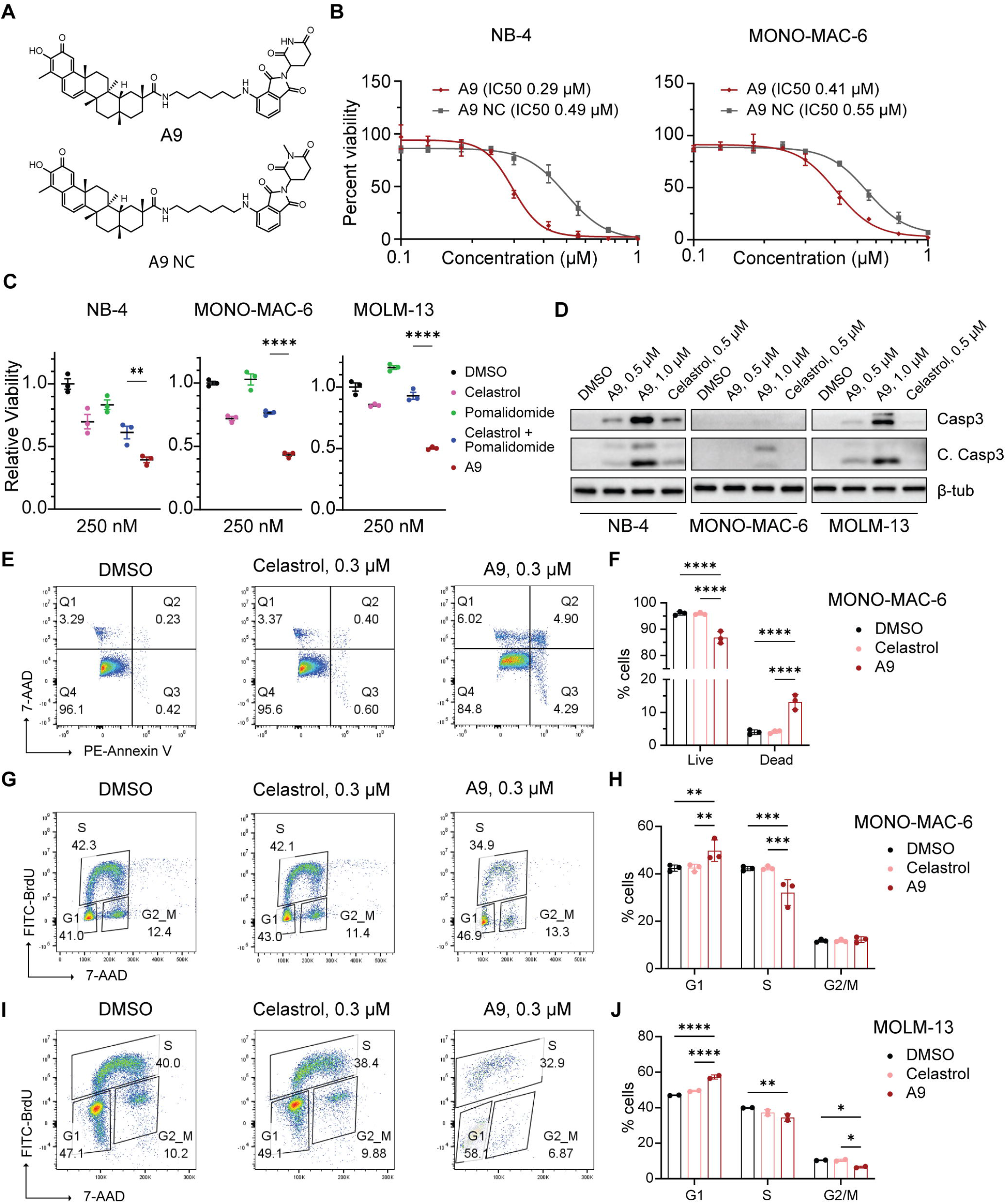
A9 induces AML cell death and growth arrest, and functions through a fundamental PROTAC mechanism. **(A)** Molecular structures of A9 and its negative control (NC) with a mutated CRBN ligand. **(B)** Kill curves of A9 vs A9 NC in NB-4 and MONO-MAC-6 cells. Cells were treated with increasing doses of A9 or N9 NC for 48 h to compute relative viabilities using MTS assay. Respective IC50 values shown within the parentheses. **(C)** Relative viabilities of NB-4, MONO-MAC-6, and MOLM-13 cells (n = 3) treated with DMSO, celastrol, pomalidomide, celastrol + pomalidomide, or A9 at 250 nM for 48 h. **(D)** NB-4, MONO-MAC-6, and MOLM-13 cells were treated with specified concentrations of A9 or celastrol to analyze caspase 3 and cleaved caspase 3 protein levels through Western Blotting with β-tubulin as a loading control. **(E–F)** MONO-MAC-6 cells were treated with DMSO, 0.3 µM celastrol, or 0.3 µM A9 for 48 hours and subjected to Annexin V/7-AAD labeling. Percentages of live (Q4) and dead (Q1 + Q2 + Q3) analyzed by flow cytometry in each condition are depicted by representative images in **E**. Cumulative data (for n = 3) summarized in a bar graph in **F. (G–J)** BrdU cell proliferation assay was performed on MONO-MAC-6 **(G–H)** and MOLM-13 **(I–J)** cells treated for 48 h with DMSO, celastrol, and A9 at given concentrations. Representative cell cycle distributions for various treatment conditions shown in **G** and **I,** and cumulative data summarized in **H** and **J**. n = 3 per treatment condition in MONO-MAC-6 or MOLM-13 cells. Data represent mean +/- SD. Statistical significance determined using one-way ANOVA followed by Fisher’s LSD test to compare specific treatment groups of interest in **C**. Two-way ANOVA followed by Tukey’s multiple comparison test was used in **F**, **H**, and **I**. *P < 0.05, **P < 0.01, ***P < 0.001, ****P < 0.0001. Only significant comparisons are marked.

Since A9 contains both celastrol and pomalidomide, we questioned whether its activity in AML cells could result from an additive effect of the two components. To address this, we treated AML cells with A9 or an equimolar combination of celastrol and pomalidomide (**Fig. 4C**). A9 exhibited superior AML cell killing to the combination, suggesting that its activity is not merely due to celastrol and pomalidomide’s combined effects. Moreover, the cytotoxic effects of celastrol were either comparable or lesser in magnitude than A9 NC in AML cells, which partly disentangles A9 associated PROTAC functions from celastrol’s inherent properties (**Supp. Fig. 4C–4E**). To further investigate the mechanism of A9-mediated cell death, we treated cells with varying concentrations of A9 and assessed apoptotic (cleaved caspase 3 and caspase 3) and necrotic (RIP kinase) markers (**Fig. 4D**). RIP kinase levels remained minimal even with A9 treatment (data not shown), whereas cleaved and full-length caspase 3 increased drastically in a dose-dependent manner (**Fig. 4D**). Functional assays demonstrated that 0.3 µM A9, unlike celastrol, significantly induces AML cell death and impedes proliferation (**Fig. 4E–4J**), recapitulating the biological outcomes previously observed with acute genetic NR4A1 and IKKβ depletions (**Fig. 2D–2H**). Together, these findings establish A9 as a potent, proof-of-principle PROTAC that induces AML cell death and cell cycle arrest through a canonical PROTAC mechanism.

### A9 induces IKK**β** and/or NR4A1 degradation through ubiquitin-proteasome pathway

At the molecular level, celastrol-based PROTACs have been shown to degrade IKKβ, NR4A1, and other proteins such as PI3Kα and CHK-1 in different non-AML contexts [19, 20]. However, a PROTAC’s degradation profile can be cell type-dependent and influenced by relative abundance of the POI, the E3 ligase, and the linker changing the structure of the ternary complex.

We treated NB-4, MONO-MAC-6, and MOLM-13 cells with different concentrations of A9 or celastrol. A9 treatment led to degradation of both IKKβ and NR4A1 in a dose-dependent manner (with nanomolar DC_50_) as shown by Western Blotting (**Fig. 5A–5B** and **Supp Fig. 5A showing summarized data**). Flow cytometry further confirmed reduced NR4A1 and IKKβ levels in A9-treated AML cells (**Supp Fig. 5B–5D**). We found that A9 can also degrade IKKβ and NR4A1 in KMT2A::MLLT3 cells, a mouse AML model commonly used for in vivo studies (**Supp Fig. 5E**). PI3K and CHK-1—two other reported targets for celastrol—were either poorly expressed or not consistently degraded relative to DMSO or celastrol-treated controls (**Supp. Fig. 5F–5H).**

**Figure 5:**
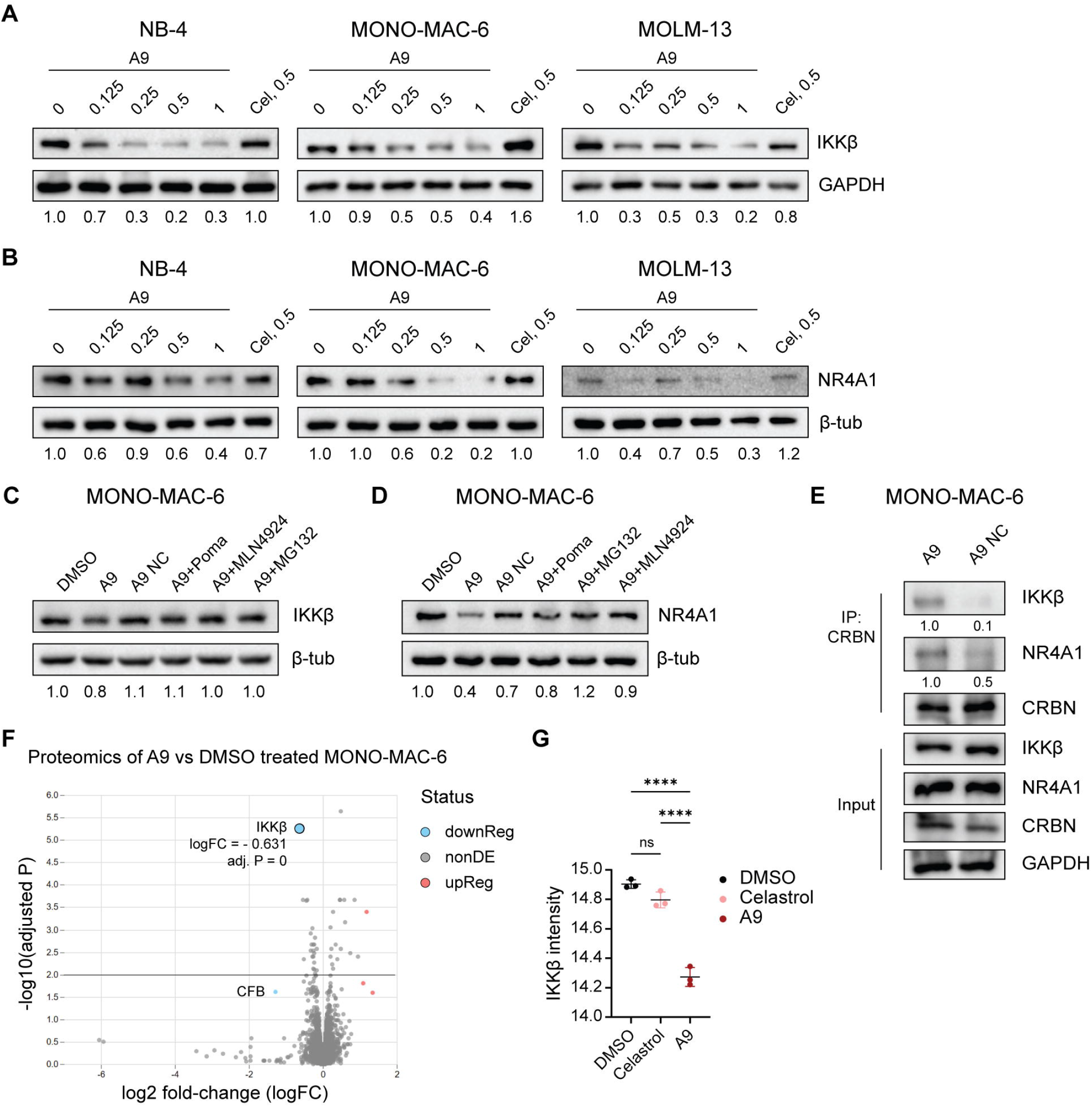
A9 induces IKKβ and NR4A1 degradation through ubiquitin-proteasome pathway. **(A– B)** NB-4, MONO-MAC-6, and MOLM-13 cells were treated with increasing concentrations of A9 or 0.5 µM celastrol and Western Blotting analyses of IKKβ and NR4A1 (using GAPDH or β-tubulin as loading controls) were performed on the cell lysates. **(C)** MONO-MAC-6 cells were treated with DMSO; 0.25 µM A9 alone or after 1 h inhibitor pretreatment (5 µM pomalidomide, 0.25 µM MG132, or 0.5 µM MLN4924); or just 0.25 µM A9 NC for 16 h. Western Blotting was performed to analyze the IKKβ protein levels with respect to the β-tubulin loading control. **(D)** MONO-MAC-6 cells were treated with DMSO; 0.5 µM A9 alone or 1 h inhibitor pretreatment (5 µM pomalidomide, 0.25 µM MG132, or 0.5 µM MLN4924); or just 0.5 µM A9 NC for 16 h. Western Blotting was performed to analyze the NR4A1 protein levels with respect to the β- tubulin loading control. **(E)** MONO-MAC-6 cells were initially treated for 1 h with 5 µM MG132 followed by co-treatment of 10 µM A9 or A9 NC for additional 3 h. Immunoprecipitates of anti- CRBN antibody were immunoblotted to test the co-immunoprecipitation (co-IP) of IKKβ and NR4A1. **(F)** Volcano plot providing an overview of the differentially regulated proteins revealed by the proteomic analyses of MONO-MAC-6 cells treated for 16 h with DMSO or 0.5 µM A9 (n = 3 per condition). Two most highly downregulated proteins marked on the plot. **(G)** Log2 VSN (Variance Stabilizing Normalization) normalized exclusive intensities of the IKKβ protein in DMSO, celastrol, and A9 treated cells quantified by proteomic analysis. Data are presented as mean +/- SD. One-way ANOVA with Tukey’s multiple comparison test used for statistical analysis in **G**. Numbers below the western Blotting images indicate relative pixel densities of the IKKβ or NR4A1 protein levels normalized to corresponding loading controls. ****P < 0.0001.

To assess involvement of the ubiquitin-proteasome system (UPS) in IKKβ and NR4A1 degradation, we examined whether inhibitors of the pathway can attenuate A9-induced target degradation. We found that co-treatment of competitive E3 ligase inhibitor (pomalidomide), proteasome inhibitor (MG132), or neddylation inhibitor (MLN4924) can partially or completely reverse A9-mediated IKKβ and NR4A1 degradation in MONO-MAC-6 cells (**Fig. 5C–5D**). A9 NC, that cannot bind to CRBN, also served as a control in these experiments. Similar results were also observed in A375 melanoma cells, suggesting that A9-induced IKKβ and NR4A1 degradation via CRBN and UPS engagement is not limited to AML (**Supp. Fig. 5I–5J**).

Ternary complex formation between PROTAC, protein target, and E3 ligase is a prerequisite for subsequent target degradation. We observed co-immunoprecipitation of endogenous IKKβ and NR4A1 with CRBN in A9 treated MONO-MAC-6 cells, confirming A9-induced ternary complex formation (**Fig. 5E**). Such co-immunoprecipitations were also demonstrated in HEK293T cells using exogenously expressed Flag-CRBN and Flag-NR4A1 (**Supp. Fig. 5K–5L**). To gain a broader insight into the full spectrum of proteins degraded by A9 as well as distinguish its degradation profile from celastrol’s intrinsic pharmacology (in the context that celastrol has been reported to have multiple E3 ligase engagements [49]), an unbiased proteomic analysis was performed on A9 treated MONO-MAC-6 cells with DMSO and celastrol treated cells serving as controls. We found that A9 and celastrol exhibit strikingly different degradation profiles when compared to DMSO controls (**Fig. 5F and Supp. Fig. 5M**). Of interest, IKKβ was the second most downregulated protein with the highest statistical significance in A9 treated cells while its levels remained unchanged in celastrol treated cells (**Fig. 5G**). Although CFB exhibited highest degree of downregulation with statistical significance (p = 0.03), its intensities across all the samples were close to the minimal detection threshold of the technique and was detected only in 1 out of 3 DMSO, 2 out of 3 A9, and none of the celastrol treated samples. NR4A1 was, however, not detected in any samples. Taken together, our results demonstrate that A9 is a potent PROTAC degrader of IKKβ and NR4A1 with pharmacological effects distinct from that of its celastrol warhead.

### A9-induced AML cytotoxicity is dependent on the degradation of IKK**β** and/or NR4A1

To determine whether degradation of IKKβ contributes to A9-induced AML suppression, we overexpressed lysine mutant IKKβ—carrying K147R, K301R, K418R, K555R, and K703, and resistant to ubiquitination and proteasomal degradation—in MONO-MAC-6 cells. We found that expression of degradation resistant IKKβ in AML cells make them less sensitive to A9 cytotoxicity (**Fig. 6A**). Corresponding Western Blotting results demonstrated that while endogenous WT IKKβ and NR4A1 underwent degradation due to A9 treatment both in EV and mutant cells, A9 stabilized the mutant IKKβ in a dose dependent manner, putatively why the cells become less sensitized to A9 cytotoxicity (**Fig. 6B**). Furthermore, *IKBKB* silenced MONO- MAC-6 cells also were resistant to A9 at 0.25 µM and 0.5 µM as opposed to their EV counterparts (**Fig. 6C–6D**), as well as NR4A1-silenced cells (**Fig. 6E–6F**). In subsequent experiments with varying depletions of NR4A1 and IKKβ in KD cells, we have also observed partial resistance at above concentrations with the overall trend remaining unchanged (data not shown). These data are a bit counter-intuitive; however, our Western Blotting results show that *IKBKB* KD also leads to pronounced downregulation of NR4A1 and vice versa (**Fig. 2B**), and further target degradation by A9 was only minimal in such contexts (**Fig. 6D** and **Fig. 6F**). The loss of both targets in the single gene KD cells (with compensatory mechanisms already in place) is presumably why they become resistant to A9 induced cytotoxicity in tested experimental conditions. We also observed a similar trend of partial or complete resistance to A9 killing in *IKBKB* KO MONO-MAC-6 cells (that exhibit reduced proliferation compared to EV, **Supp. Fig. 6A–6D**) and *NR4A1* KO MOLM-13 cells (**Fig. 1F** and **Supp. Fig. 6E**).

**Figure 6:**
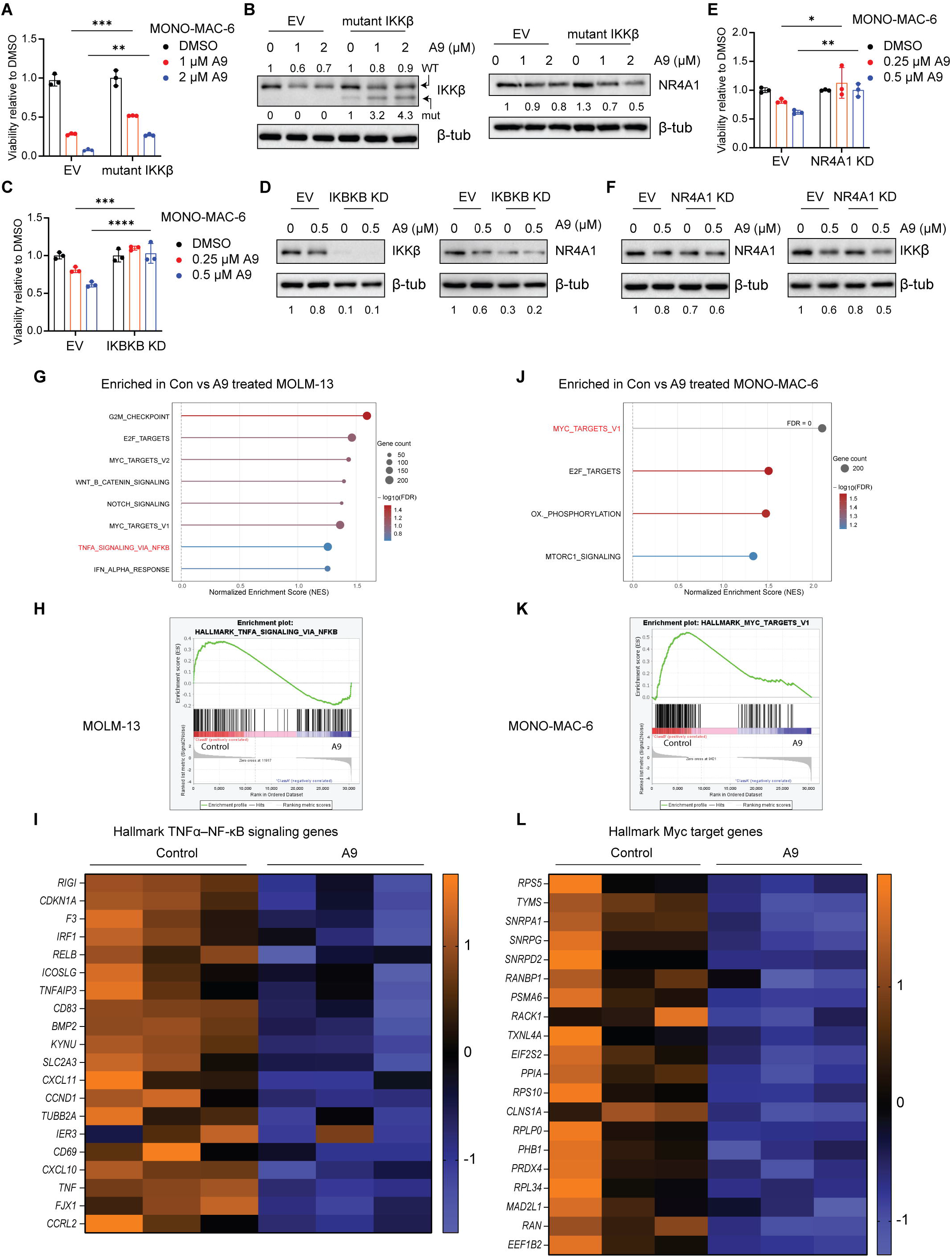
A9 reduces AML cell viability in an IKKβ- and/or NR4A1-dependent manner. **(A)** MONO-MAC-6 cells were transduced with EV lentiviral vector or that expressing lysine-mutant IKKβ (sequence available in supplementary information). EV and mutant IKKβ expressing cells (n = 3) were treated (7 days after terminating lentiviral infection) with DMSO, 1 µM, or 2 µM A9 for 48 h prior to cell viability assessment via MTS assay. Bar diagram depict viabilities of A9 treated EV and mutant cells relative to their corresponding DMSO treated counterparts to compare their sensitivities to A9. **(B)** Western Blotting images showing the WT and mutant IKKβ as well as NR4A1 protein levels in transduced MONO-MAC-6 cells following their treatment with DMSO or A9 for 16 h. **(C)** EV and *IKBKB* KD cells (n = 3 for EV and KD) were treated (7 days after terminating lentiviral infection) with DMSO, 0.25 µM, or 0.5 µM A9 for 48 h prior to cell viability assessment via MTS assay. Bar diagram depict viabilities of A9 treated EV and KD cells relative to their corresponding DMSO counterparts to compare their sensitivities to A9. **(D)** Western Blotting images showing the IKKβ and NR4A1 protein levels in EV and *IKBKB* KD MONO-MAC-6 cells after treating for 16 h with DMSO or 0.5 µM A9. **(E)** EV and *NR4A1* KD cells (n = 3) were treated (7 days after terminating lentiviral infection) with DMSO, 0.25 µM, or 0.5 µM A9 for 48 h prior to cell viability assessment via MTS assay. Bar diagram depict viabilities of A9 treated EV and KD cells relative to their corresponding DMSO treated counterparts to compare their sensitivities to A9. **(F)** Western Blotting images showing the levels of NR4A1 and IKKβ in EV and *NR4A1* KD MONO-MAC-6 cells following their treatment with DMSO or 0.5 A9 for 16 h. **(G)** MOLM-13 cells (n = 3) were treated with DMSO or 0.125 µM A9 for 72 h, and bulk RNA-sequencing and analysis were performed. Dot plot of significantly enriched pathways in GSEA analysis of DMSO control- vs A9-treated MOLM-13 samples. **(H)** Enrichment plot showing enhanced TNFα signaling via NF-κB in control MOLM-13 cells compared to the A9-treated. **(I)** Heatmap of top 20 genes, belonging to the TNFα–NF-κB signaling pathway, upregulated in control- vs A9-treated MOLM-13 cells. **(J)** MONO-MAC-6 cells (n = 3) were treated with DMSO or 0.25 µM A9 for 72 h, and bulk RNA-sequencing and analysis were performed. Dot plot of significantly enriched pathways in DMSO control- vs A9-treated MONO-MAC-6 cells. **(K)** Enrichment plot showing enriched Myc target genes in control MONO- MAC-6 cells compared to the A9-treated. **(L)** Heatmap of top 20 differentially expressed Myc target genes in control- vs A9-treated MONO-MAC-6 cells. Data represent mean +/- SD. Two- way ANOVA followed by Tukey’s multiple comparison test was used to compare groups of interest in **A**, **C**, **E**. **\***P < 0.05, **P < 0.01, ***P < 0.001, ****P < 0.0001. Numbers below the western blotting images indicate relative pixel densities of the IKKβ or NR4A1 protein levels normalized to corresponding loading controls.

To understand downstream pathways altered by A9-mediated target degradation, we performed bulk RNA-sequencing on MOLM-13 and MONO-MAC-6 cells treated with DMSO or A9 for 72 h. GSEA analysis of differentially expressed genes (DEGs) revealed enrichment of many oncogenic pathways in control MOLM-13 cells and MONO-MAC-6 compared to their A9- treated counterparts (**Fig. 6G–6L, Supp. Fig. 6F–6I**). Notably, the TNFα–NF-kB signaling pathway was significantly enriched in control- vs A9-treated as well as some other cancer pathways (Notch, Wnt-β-catenin), which were also enriched in NR4A1 Hi (vs Lo) AMLs in the TCGA dataset (**Supp. Fig. 2C**). These shared pathway alterations, including the downregulation of NF-kB signaling, corroborate the idea that A9-induced biological changes could be largely mediated by IKKβ and/or NR4A1 degradation.

### A9 attenuated AML progression and did not induce neutrophilia in mice

To test *in vivo* efficacy of A9 against AML progression, murine KMT2A::MLLT3 cells were transplanted into sub-lethally irradiated C57BL6/J mice. The oncogenic KMT2A::MLLT3 fusion protein, resulting from the t(9;11)(p22;q23) translocation, is commonly observed in AML patients [50, 51]. A9-treated mice exhibited slower AML progression, demonstrated by reduced blood AML cell counts compared to the vehicle-treated cohort (**Fig. 7A**).

**Figure 7:**
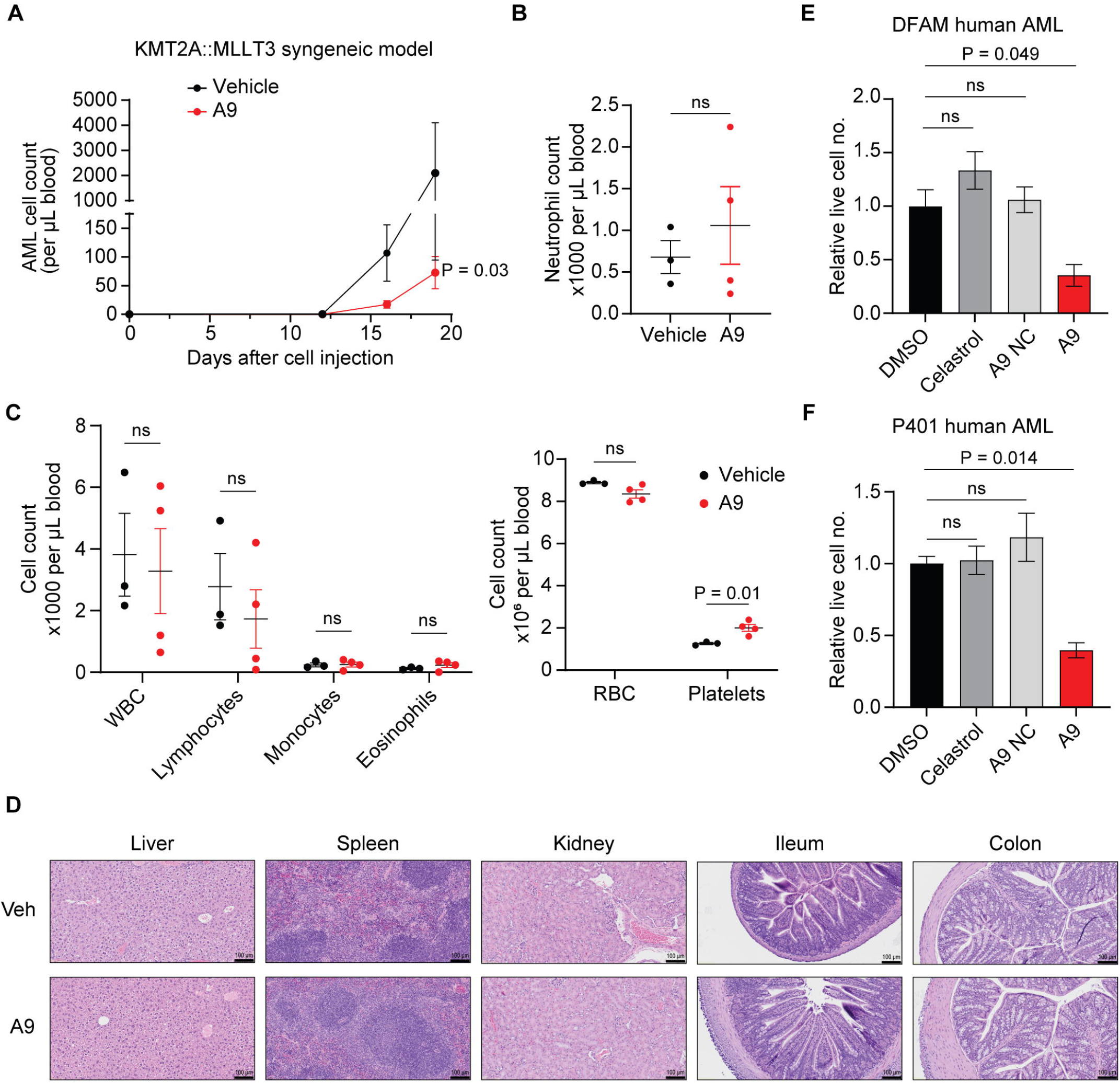
A9 attenuated AML progression and did not induce neutrophilia in mice. **(A)** C57BL6/J mice were irradiated with 6.5 Gy X-ray and 2x10^4^ syngeneic KMT2A::MLLT3 (aka MLL-AF9) AML (GFP tagged) cells were transplanted into each mouse via tail vein injection. A9 formulation (at 4 mg/kg dosage) or the vehicle were administered into the mice (n = 4 per group) on the day after cell injection and this was repeated every 3 days. The A9 dose was reduced to 2 mg/kg after the first four administrations. AML cell count in mouse blood was periodically quantified by flow cytometry to generate the growth curves shown. Statistical significance determined using two-way ANOVA. **(B–C)** Healthy C57BL6/J mice were administered with A9 drug or vehicle following the same treatment regimen (first four doses of 4 mg/kg every 3 days reduced to 2 mg/kg thereafter) as in Fig. A. Results of complete blood count study of vehicle (n = 3) and A9 (n = 4) treated mice on are shown. Statistical significance determined by student’s T-test. **(D)** Representative pictures of H&E-stained tissue sections harvested from the vehicle and A9-treated C57BL6/J mice described above. (**E–F**) Human AML cells were treated with 250 nM celastrol, A9 NC, or A9 for 24 h and the number of live cells were manually counted by trypan blue exclusion method and normalized to the DMSO control. Statistical significance determined using one-way ANOVA with Tukey’s multiple comparison test. Data represent mean +/- SEM.

The best-known adverse effect of IKKβ inhibition via small molecule inhibitors is induction of neutrophilia and subsequent widespread inflammation [10, 11]. To assess this risk, healthy C57BL6/J mice were treated with A9 or vehicle, and CBC was performed. Importantly, we found no significant difference in the neutrophil count in their blood (**Fig. 7B**), likely due to lowest expression of CRBN in neutrophils among all human immune cell types (**Supp. Fig. 7**). Counts of other cell types, including WBCs, lymphocytes, monocytes, and eosinophils, also displayed no significant difference (**Fig. 7C, left panel**), with slightly increased platelets (**Fig, 7C, right panel**). Moreover, H&E-stained tissue sections exhibit no obvious abnormalities in A9- treated mice (**Fig. 7D**).

We tested the efficacy of A9 in two human AML specimens (DFAM68555 and P401) with celastrol and A9 NC as controls (**Fig. 7E–7F**). A9 drastically reduced the number of viable cells, whereas cell viability in DMSO, celastrol, and A9 NC conditions were similar. Overall, A9 demonstrated strong anti-AML activity in clinically relevant disease models without inducing life-threatening hematological abnormalities *in vivo*.

## Discussion

Here we show that a single PROTAC drug can effectively degrade IKKβ and/or NR4A1, apparently leading to robust AML suppression *in vitro* and *in vivo*. The CRBN-based A9 PROTAC evades adverse hematological effects that have hindered the development of IKKβ small molecule inhibitors.

In our analysis of a published scRNA-seq dataset from AML patients [44], we observed a strong correlation between *IKBKB* expression and enrichment of proinflammatory signatures, which connotes poor clinical outcomes, in CD34^+^ leukemic BM cells [44]. We found that IKKβ is particularly required for AML cell proliferation in vitro. The proof-of-concept PROTAC, A9, is a potent degrader of IKKβ and can suppress AML pathogenesis in an IKKβ-dependent manner. Provided the AML cell line we employed (i.e., MONO-MAC-6) for IKKβ functional studies is of (pro)monocytic origin, our results are consistent with prior studies that have deciphered the oncogenic role of inflammatory pathways (particularly NF-κB signaling) in monocytic AMLs [52]. Future studies in our lab will more comprehensively study the role of IKKβ as well as NR4A1 in AML models of different myeloid cell lineages and differentiation status.

A9 also degraded NR4A1, another important AML target, which potentially contributes to A9-induced AML cell death. Our analysis of TCGA-AML datasets found that AML patients exhibit varying levels of NR4A1 expression, and that high expression is associated with poor prognosis and shorter overall survival. Our genetic loss-of-function studies show that NR4A1 depletion primarily promotes cell death and differentiation, while moderately reducing cell proliferation *in vitro*; consistent with suppression of AML progression *in vivo*. Provided NR4A1 and IKKβ play non-redundant oncogenic roles in AML pathogenesis, targeting of IKKβ and/or NR4A1 could be an effective AML therapeutic strategy.

Celastrol is a promiscuous natural product that putatively binds to several proteins. Celastrol-based PROTACs have also been shown to degrade IKKβ, NR4A1, CHK1, PI3Kα, and a few others, including PP2A and OGA, in different non-AML contexts [19]. We demonstrate that A9 leads to pronounced, dose-dependent degradation of IKKβ and NR4A1 in AML cells, predominantly contributing to A9-induced cytotoxicity. Our unbiased proteomic analysis reinforces these findings by revealing IKKβ as the most significantly downregulated protein in A9, but not in celastrol, treated cells. Moreover, A9 and celastrol exhibited distinct degradation profiles, that help to disentangle A9’s superior AML suppressive effects from the intrinsic pharmacology of celastrol.

NR4A1 could play a dichotomous role in AML pathogenesis. Previous literature demonstrated that NR4A1 plays a tumor suppressive role in certain *in vitro* and mouse AML models. They showed that combined genetic ablation of *Nr4a1* and *Nr4a3* precipitates an AML-like disease in mice, although loss of either gene alone is insufficient to drive the phenotype [40]. In addition, overexpression of NR4A1 in Kasumi-1 AML cells with poor NR4A1 expression inhibits their proliferation [41]. In sum, the biphasic role of NR4A1 in AML could be primarily disease subtype-dependent (considering AML is a very heterogenous disease) and will be interesting to further investigate in the future.

Nearly half of the AML patients in the TCGA-AML dataset belonging to the poor cytogenetic risk category exhibited high expression of both IKKβ and NR4A1. The potential targeting of IKKβ and/or NR4A1 using one PROTAC could therefore benefit patients with refractory AMLs. Rather than two independent celastrol targets, NR4A1 and IKKβ (or NF-κB signaling) have been reported by many studies to closely crosstalk, modulating each other’s activity [53]. Consistently, we observed reciprocal positive regulation between IKKβ and NR4A1 in certain AML cells but not in all AML cells. Transcription of NR4A1 and other *NR4A* family genes is induced by the activation of NF-κB signaling in response to diverse inflammatory stimuli in myeloid cells [54, 55]. NR4A1 can negatively or positively feedback on the NF-κB pathway to reduce or reinforce the inflammatory response [53, 55–58]. NR4A1, for instance, has been shown to directly interact with p65 to inhibit NF-κB/p65 binding to target gene promoters, which is countered by the LPS-activated p38-a phosphorylation of NR4A1 [59]. Alternatively, NR4A1 can promote proinflammatory functions [56, 57], most notably by directly transactivating IKKi/IKKε, key components of NF-κB [56]. We found that IKKβ and NR4A1 play non-redundant oncogenic functions in AML pathophysiology, presumably why combined degradation of both NR4A1 and IKKβ by A9 induces remarkable AML suppression.

A key advantage of PROTAC therapy is the ability to degrade POIs in a tissue-selective manner, facilitated by differential E3 ligase expression, which helps avoid many adverse side effects [13, 15]. Systemic IKKβ inhibition by small molecule inhibitors is known to induce neutrophilia and widespread inflammation [10, 11]. Our data demonstrate that A9 degrades NR4A1 and IKKβ by employing the CRBN E3 ligase, which is nearly undetectable in neutrophils. Consistently, A9 treatment in mice did not cause neutrophilia or other hematological adverse effects. As such, our PROTAC design holds promise for further investigations and development in AML therapy. While we did not observe any significant alterations in the mouse PBMC counts after a short A9 treatment, further experiments focusing on normal bone marrow HSCs are required to advance clinical development of the drug.

Overall, our results reveal PROTAC-mediated degradation of IKKβ and/or NR4A1 as a novel strategy for effective AML therapy that could benefit patients with intractable AMLs.

## Data availability

All original data are available upon request. We include new single-cell RNA sequencing analysis from a publicly available dataset, GSE185381. Our bulk RNA sequencing dataset is deposited as GSE324608. TCGA datasets were downloaded for the TCGA website or UCSC Xena Browser as indicated within the corresponding sections.

## Code availability

The analyses in this study were performed using standard and publicly available R packages as indicated in the corresponding method sections for sequencing analyses. No custom code essential for reproducing the results was developed.

## Author Contributions

Study conceptualization and design: C.K.M., Y.L., Y.X., G.Z. and W.Z. Experimentation, materials, data acquisition and analysis: C.K.M., Y.L., Y.X., B.R.P., T.H.M., L.W., M.C.K, Z.J., S.A., T.I.T., V.M.S., C.Z., S.A., A.M.S., Z.Q., and J.L. Supervision and scientific interpretation of data: C.K.M., R.K., G.Z., and W.Z. Writing – original draft: C.K.M., and Y.L. Writing – review & editing: R.K., G.Z., and W.Z. with minor editing from all authors. Funding Acquisition: W.Z. All authors have read and agreed to the published version of the manuscript.

## Acknowledgments

P401 AML cells were utilized here (courtesy of Alexandra M. Stevens, Texas Children’s Hospital) and supported by grants from the Texas Children’s Hospital Pediatric Pilot Research Fund (AMS), Leukemia and Lymphoma Society (AMS), Target Pediatric AML/the Children’s Oncology Group Foundation (AMS), and gifts of funding from the Turn it Gold Fund (AMS). We thank Dr. Chen Zhao (CWRU) for sharing KMT2A::MLLT3 cells and DFAM68555 primary AML cells; Dr. Pradeep Ramalingam (Jason Butler Lab, UF) for his helpful suggestions on working with the KMT2A::MLLT3 mouse AML model and Dr. Jason Butler Lab for help with CBC analysis.

The work is partly supported by DOD/CDMRP BC200100 (PIs: W. Zhang and G. Zheng), NIH/NCI: CA269661 (W. Zhang), CA260239 (W. Zhang; D. Zhou; G. Zheng), and CA290792 (W. Zhang; G. Zheng; K.S.M. Smalley), University of Florida Health Cancer Institute grant P30CA247796 (PI: T. George), an endowment fund from the Dr. and Mrs. James Robert Spenser Family (W. Zhang).

