## Supplementary Information for "PROTAC-Mediated Targeting of IKKβ and NR4A1 for AML Therapy"

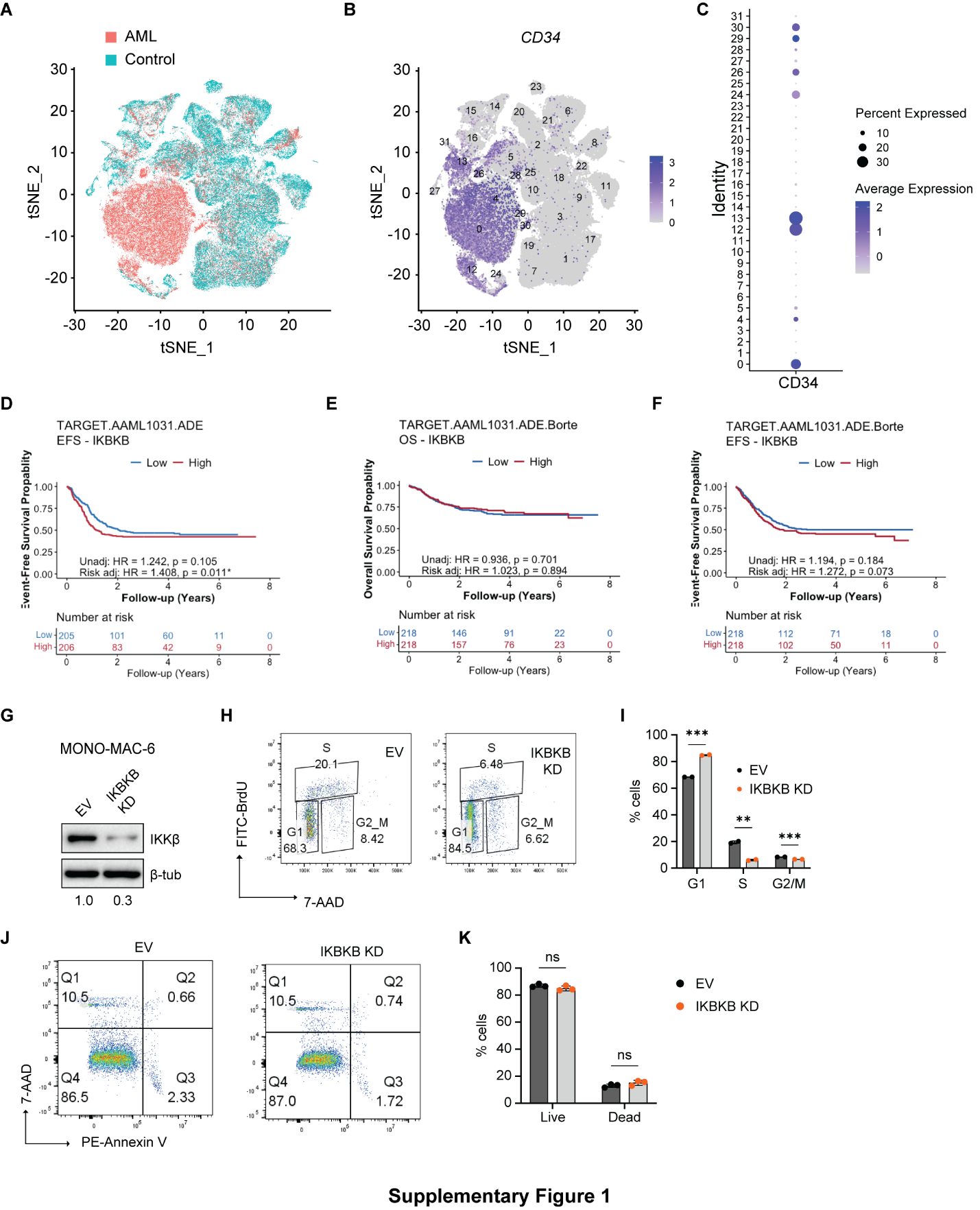

**Supplementary Figure 1: (A)** Feature plot depicting single-cell tSNE for each group. BM cells from AML patients and healthy controls have different global structures of RNA expression. Feature **(B)** and dot **(C)** plots depicting *CD34* gene expression on single-cell clusters from patients with AML and healthy controls. Clusters 0, 4, 12, 13, 24, 26, 29, and 30 were considered positive for *CD34* gene expression and applied to correlation analysis in **Fig. 1B**. **(D)** Kaplan-Meier plot comparing the event free survival (EFS) of *IKBKB* high vs low patients (separated by median) in the ADE (cytarabine, daunorubicin, and etoposide) treatment arm of the Target-AAML1031 clinical trial patient cohort. **(E–F)** Kaplan-Meier plots comparing the event overall survival (OS, in **E**) and event free survival (EFS, in **F**) of *IKBKB* high vs low patients in the ADE + Bortezomib treatment arm of the Target-AAML1031 clinical trial patient cohort. **(G)** Western Blotting images showing IKKβ levels in EV control vs *IKBKB* knockdown (KD) MONO-MAC-6 cells generated by transducing corresponding lentiviral shRNAs. **(H–I)** Empty vector (EV) and *IKBKB* KD MONO-MAC-6 cells were incubated with BrdU for 2 h to perform cell proliferation assay via anti-BrdU antibody and 7-AAD labeling. Representative images show the distribution of cells in different phases of the cell cycle in **H** and cumulative data summarized in a bar graph in **I**. n = 2 (EV and *IKBKB* KD). **(J)** Annexin V/7-AAD labeling of EV and *IKBKB* KD cells reveal the percentages of live (Q4), apoptotic (late, Q2 + early, Q3), and necrotic (Q1) cell populations (5 days post termination of lentiviral infection) in the flow diagrams. **(K)** Cumulative data from annexin V/7-AAD labeling (for n = 3 biological repeats) depicting the percentage of live and dead (Q1+Q2+Q3) populations. Unadjusted and risk group–adjusted hazard ratios were obtained using Cox proportional hazard models in **D**, **E**, and **F**. Data are presented as mean +/- SD in **I** and **K**. Unpaired two-tailed T-tests were used for statistical analyses in **I** and **K**.

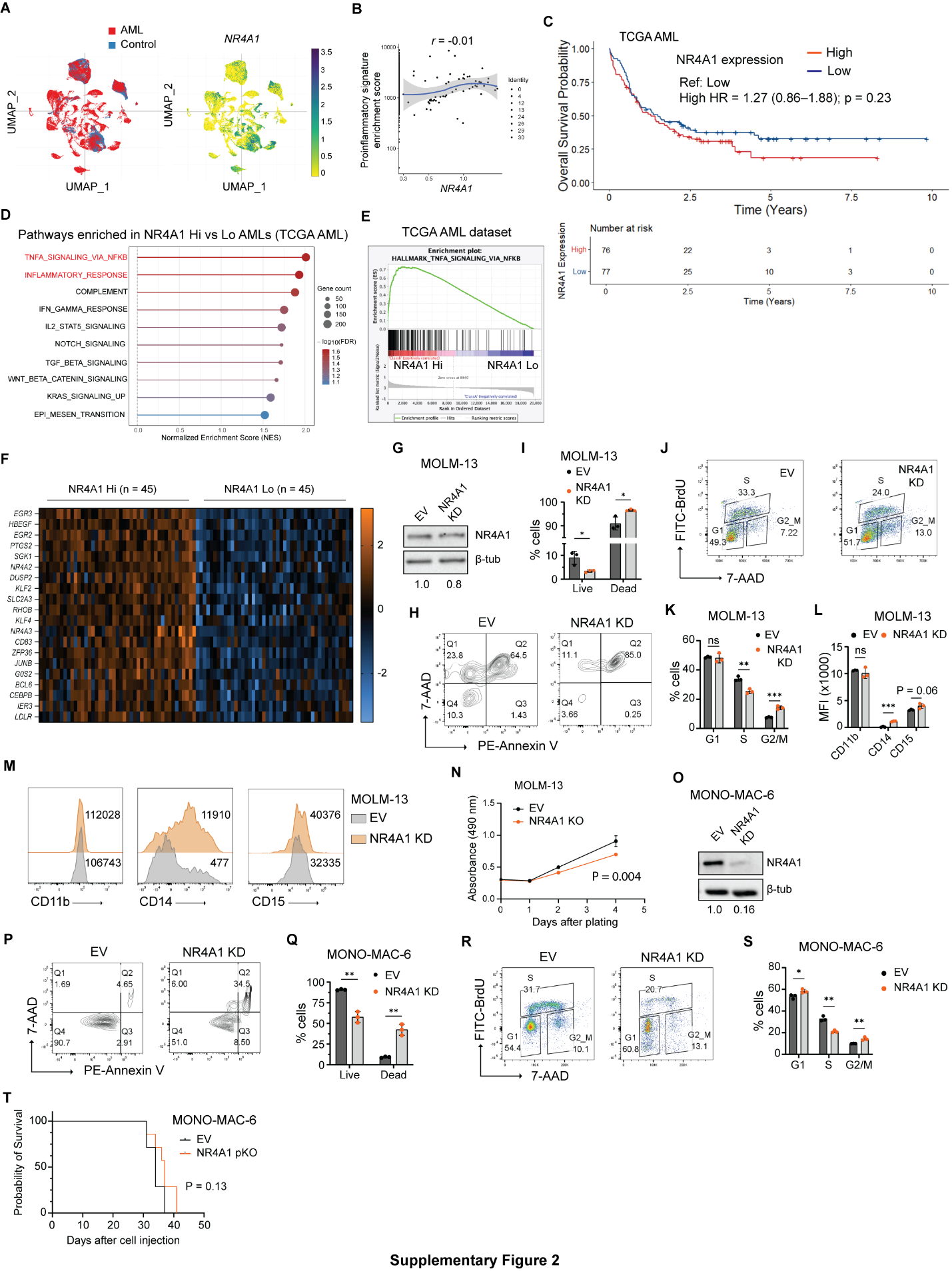

**Supplementary Figure 2: (A)** UMAP plots displaying relative *NR4A1* expression in various AML and healthy control BM cell clusters. Data downloaded from the interactive Single Cell Portal gene expression dataset deposited by Lasry et al. [1]. **(B)** Feature scatter plot depicting no correlation between *NR4A1* gene expression and enrichment score of proinflammatory signature in the CD34+ cells derived from patients with AML. **(C)** Kaplan-Meier curves representing the overall survival probability vs time (in days) of NR4A1 high and NR4A1 low expressors (separated by median) in GDC TCGA-AML patient dataset. **(D)** AML patients in TCGA database were segregated into 4 quartiles based on their relative expression of *NR4A1*. Dot plot of significantly enriched pathways enriched in NR4A1 High vs NR4A1 Low patients as revealed by gene set enrichment analysis (GSEA) of hallmark (H) gene sets. **(E)** GSEA plot revealing signifiant enrichment of the TNFα­_­_–NF-κB signaling genes in NR4A1 High vs Low AMLs. **(F)** Heat map of top 20 genes upregulated in NR4A1 Hi AMLs compared to the NR4A1 Lo. **(G)** Western Blotting images showing NR4A1 levels in EV control vs *NR4A1* knockdown (KD) MOLM-13 cells generated by transducing lentiviral shRNAs. **(H–I)** Annexin V/7-AAD staining was performed on EV and *NR4A1* KD MOLM-13 cells for flow cytometric analysis of live (Q4), apoptotic (late, Q2 and early, Q3), and necrotic (Q1) cell populations. Representative images are shown in **H** and cumulative percentages of live and dead (Q1+Q2+Q3) populations in **I**. n = 3 (EV and KD). **(J–K)** EV and *NR4A1* KD MOLM-13 cells were incubated with BrdU for 2 h to perform cell proliferation assay using anti-BrdU antibody and 7-AAD labeling. Representative images show the distribution of cells in different phases of the cell cycle in **J** and cumulative data summarized in a bar graph in **K**. **(L)** Mean fluorescence intensities (MFI) of mature monocytic differentiation markers, CD11b, CD14, and CD15, measured by flow cytometry in *EV* and *NR4A1 KD* MOLM-13 cells. **(M)** Representative images for the expression of CD11b, CD14, and CD15 in EV vs *NR4A1* KD MOLM-13 cells. Number represent mean fluorescence intensities. **(N)** Cell growth curves of EV control and NR4A1 KO Molm-13 cells generated via MTS assay. All cells were plated on Day 0 and MTS reagents were added to a new set of wells each consecutive day to assess cell growth. Data represent mean +/- SD and the curves were compared by two-way ANOVA. **(O)** Western Blotting images showing NR4A1 expression levels in EV and *NR4A1* KD MONO-MAC-6 cells generated by using lentiviral shRNAs. **(P­­–Q)** Annexin V/7-AAD labeling of EV and *NR4A1* KD MONO-MAC-6 cells to assess live and dead (Q1+Q2+Q3) cell percentages. Representative flow images are shown in **P** and cumulative data in **Q**. n = 3 (EV and KD). (**R–S**) BrdU cell proliferation assay was performed on EV and *NR4A1* KD MONOMAC-6 cells after a 2 h BrdU treatment. Representative images show the distribution of cells at different phases of the cell cycle in **R** and cumulative data summarized in a bar graph in **S**. **(T)** Kaplan-Meier plot comparing the survival of EV (n = 7) vs *NR4A1* pKO (n = 7) MONO-MAC-6 injected mice. Unpaired two-tailed T-tests were performed for statistical analysis in **I, K, L, Q**, and **S**. Log rank test used for statistical analysis in **T**.

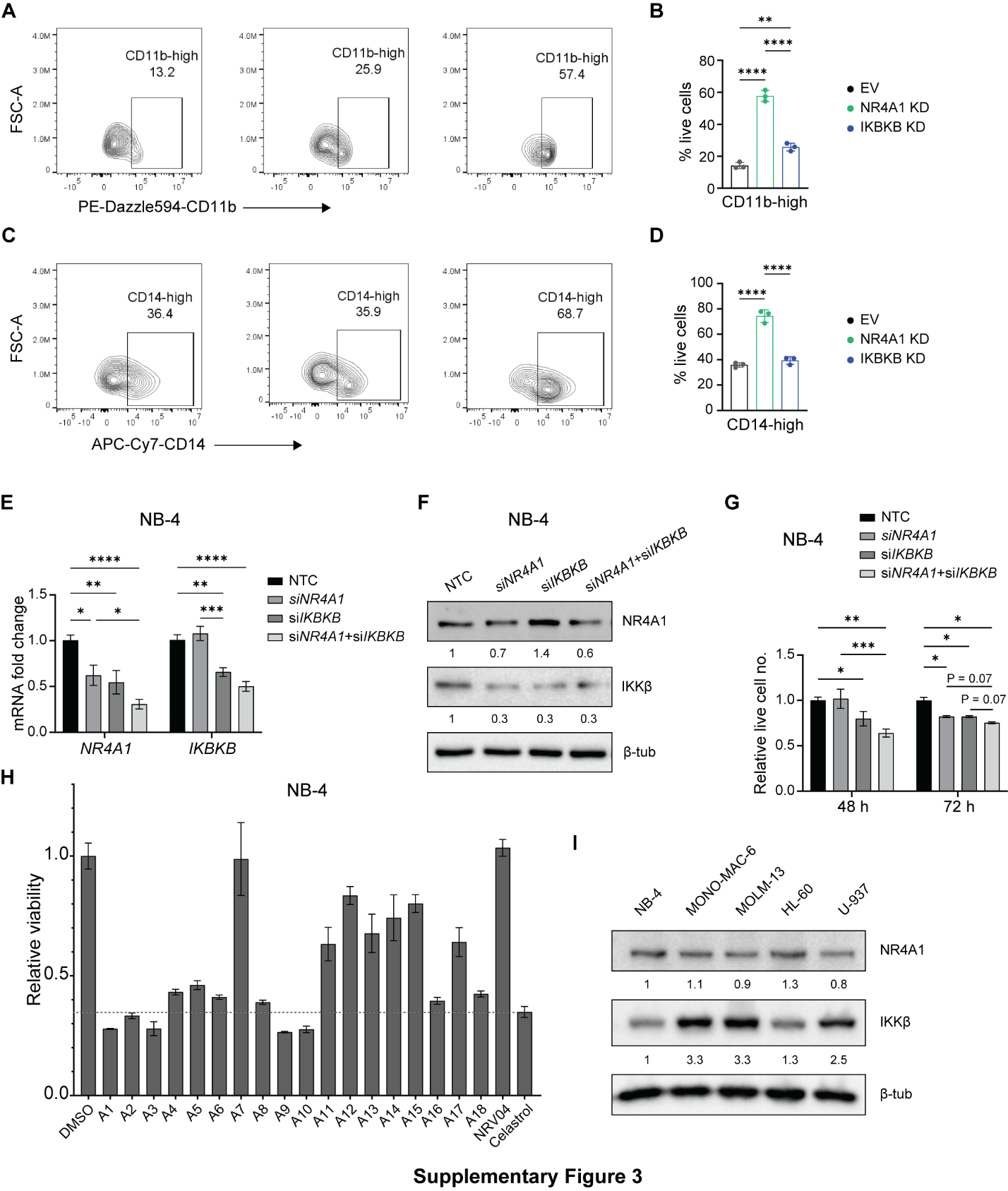

**Supplementary Figure 3:**

**(A–B)** Flow cytometric analysis of MONO-MAC-6 EV, *NR4A1* KD, *IKBKB* KD cell populations having high expression of CD11b, a myeloid differentiation marker. Representative contour plots shown in **A** and the cumulative data in **B**. **(C–D)** Flow cytometric quantification of MONO-MAC-6 EV, *NR4A1* KD, *IKBKB* KD cell populations with high expression of CD14, a mature monocytic differentiation marker. Representative contour plots shown in **C** and the cumulative data in **D**. Data represent mean +/- SD in **B** and **D** from n = 3 biological replicates and one-way ANOVA with Tukey’s multiple correction tests used for statistical analysis. **(E)** Transcript levels of *NR4A1* and *IKBKB* in NB-4 cells 48 h after the nucleofection of non-targeted control (NTC) siRNA, si*NR4A1*, si*IKBKB*, or both si*NR4A1* and si*IKBKB*. **(F)** Western blot images of NR4A1 and IKKβ with densitometric quantification of the proteins in control and various KD cells harvested 48 h after the nucleofection. **(G)** Relative live NB-4 cell counts in the above knockdown conditions 48 h and 72 h after siRNA nucleofection. Data are presented as mean +/- SEM. Statistical significance determined using two-way ANOVA followed by Fisher’s LSD test to compare specific groups of interest. **(H)** NB-4 AML cells were treated with a panel of celastrol based PROTACs or just celastrol for 48 h and their viabilities relative to DMSO control were analyzed via MTS assay. Bars represent mean +/- SD. **(I)** Western Blotting images showing variable expressions of NR4A1 and IKKβ in AML cell lines used in the current study. Numbers below indicate relative expression levels of the proteins determined via densitometric analysis in ImageJ. *P < 0.05, **P < 0.01, ***P < 0.001, ****P < 0.0001.

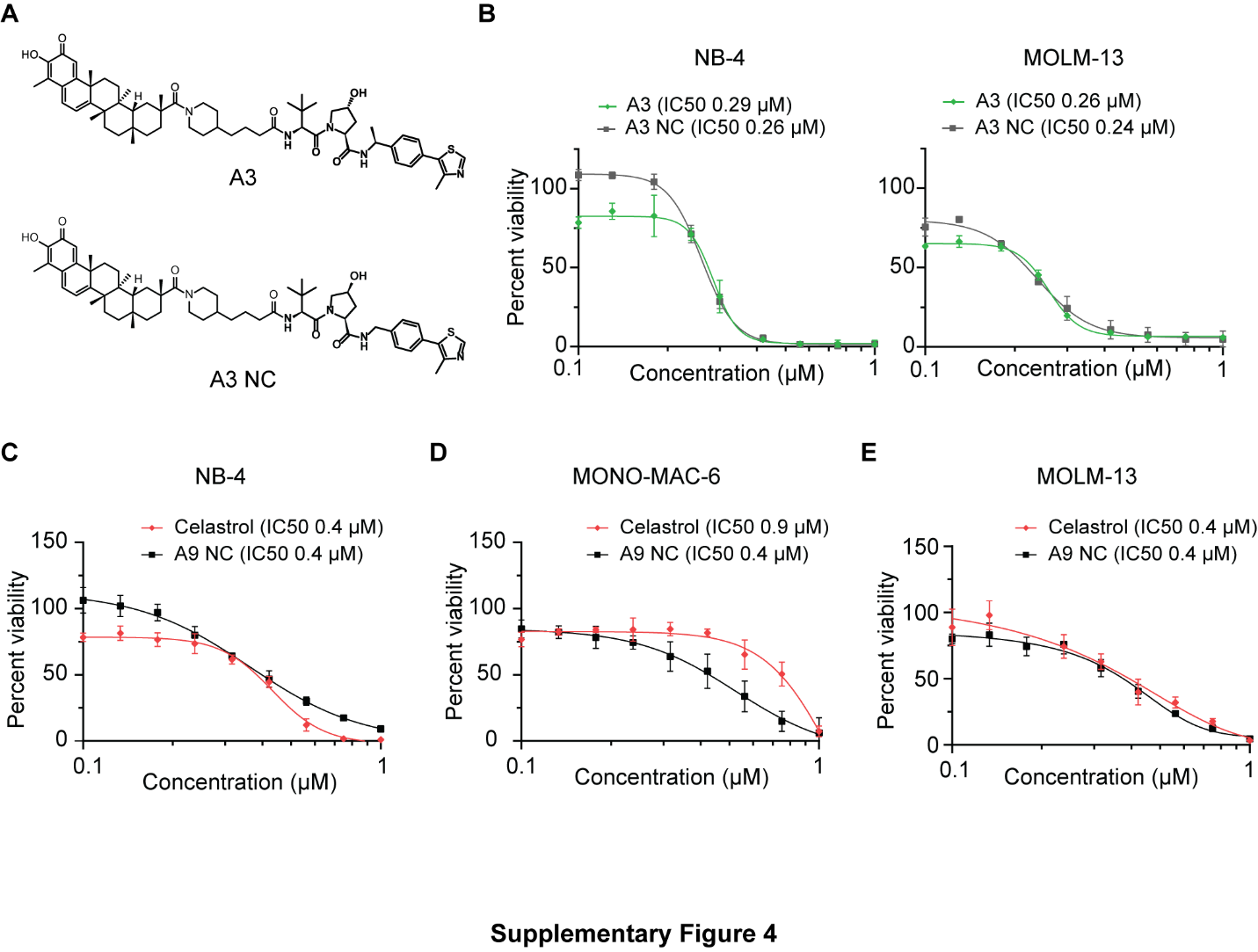
**Supplementary Figure 4:** **(A)** Molecular structures of A3 and its negative control (A3 NC) with a mutated VHL ligand. **(B)** Kill curves of A3 vs A3 NC in NB-4 and Molm-13 cells. Cells were treated with increasing doses of A3 or A3 NC for 48 h and their viabilities relative to the DMSO treatment condition were analyzed via MTS assay. **(C–E)** Comparative evaluation of celastrol vs A9 NC cytotoxicity in NB-4 **(C)**, MONO-MAC-6 **(D)**, and MOLM-13 **(E)** cells as shown by dose-response curves generated via MTS assays. Data points on the kill curves represent mean +/- SD. IC50 values indicated inside the parentheses.

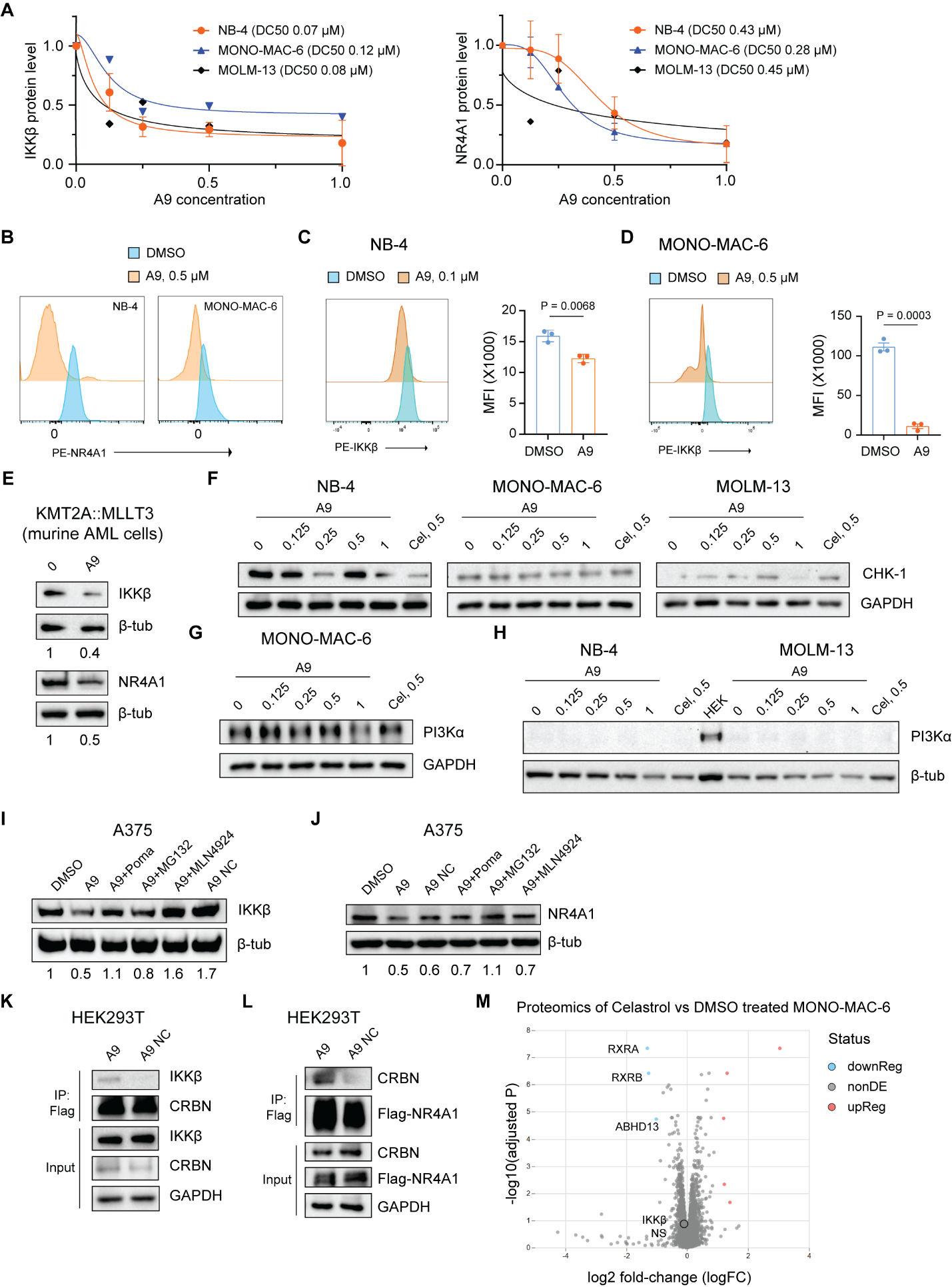

**Supplementary Figure 5: (A)** Densitometric analysis of western blots was performed to compute relative levels of the IKKβ and NR4A1 protein levels in AML cells treated with A9 in a dose dependent manner. Dose-response curves were plotted to calculate half-maximal degradation concentration (DC50) values of A9 in the indicated AML cell lines. **(B)** Flow cytometric analysis of the NR4A1 protein levels in treated with NB-4 and Monomac-6 cells after 16 h DMSO vs. 0.5 µM A9 treatment. **(C)** Flow cytometric evaluation of IKKβ levels in NB-4 cells treated with DMSO vs 0.1 µM A9 for 24 h. Representative images on the left and cumulative data for n = 3 presented in a bar diagram (right). **(D)** Flow cytometric evaluation of IKKβ levels in MONO-MAC-6 cells treated with DMSO vs 0.5 µM A9 for 24 h. Representative images on the left and cumulative data for n = 3 presented in a bar diagram (right). **(E)** Western blot results depicting IKKβ and NR4A1 levels in KMT2A::MLLT3 (murine AML) cells treated with DMSO or 0.125 µM A9. Numbers indicate relative levels of the IKKβ and NR4A1 proteins normalized to the corresponding β-tub loading controls. **(F–H)** Western blot results showing CHK-1 (**F**) and PI3Kα **(G–H)** levels in AML cells treated with A9 (0–1 µM) or celastrol (0.5 µM) for 16 h. HEK cell lysate used as a positive control for PI3Kα immunoblot in **H**. **(I)** A375 cells were treated with DMSO; 0.5 µM A9 alone or after 2 h inhibitor pretreatment (5 µM pomalidomide, 0.5 µM MG132, or 0.5 µM MLN4924); or just 0.5 µM A9 NC for 16 h. Western blot was performed to analyze the IKKβ protein levels with respect to the β-tubulin loading control. **(J)** A375 cells were treated with DMSO; 0.5 µM A9 with or without inhibitor (5 µM pomalidomide, 0.5 µM MG132, or 0.5 µM MLN4924) co-treatment; or just 0.5 µM A9 NC for 16 h. Western blot was performed to analyze the NR4A1 protein levels with respect to the β-tubulin loading control. **(K)** Flag-CRBN was transiently expressed in HEK293T cells. After initial 1 h treatment with 5 µM MG132, Flag-CRBN expressing cells were co-treated with 10 µM A9 or A9 NC for 3 h. Flag-CRBN was immunoprecipitated with anti-M2 Flag antibody and IKKβ was immunoblotted to test its co-immunoprecipitation with CRBN in A9 treated cells. **(L)** Flag-NR4A1 was transiently expressed in HEK293T cells. After initial 1 h treatment with 5 µM MG132, Flag-CRBN expressing cells were co-treated with 10 µM A9 or A9 NC for 3 h. Flag-NR4A1 was immunoprecipitated with anti-M2 Flag antibody and CRBN was immunoblotted to test its co-immunoprecipitation with NR4A1 A9-treated cells. Numbers below the western blot images indicate relative pixel densities of the IKKβ or NR4A1 protein levels normalized to corresponding loading controls. **(M)** Volcano plot providing an overview of the differentially regulated proteins revealed by the proteomic analyses of MONO-MAC-6 cells treated for 16 h with DMSO or 0.5 µM celastrol (n = 3 per condition). Data are presented as mean +/- SD in **A**, **C**, and **D**. Unpaired two-tailed T test used for statistical analysis in **C** and **D**.

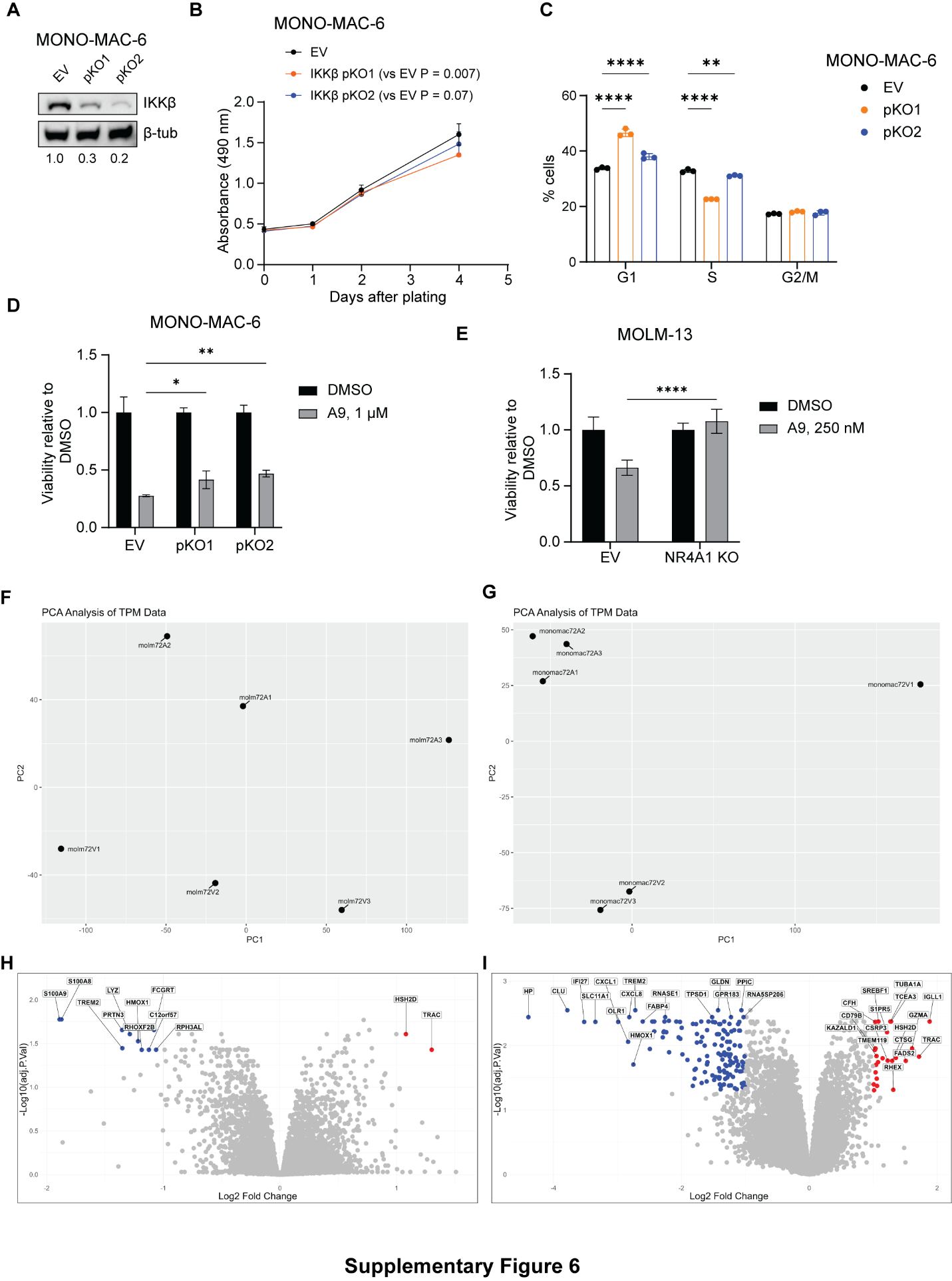

**Supplementary Figure 6: (A)** Pooled IKKβ knockout MONO-MAC-6 cells, pKO1 and pKO2, were generated via lentiCRISPRv2 technique using different guide RNAs targeting the *IKBKB* gene. Western blot images showing reduced expression of IKKβ in pKO1 and pKO2 with respect to the EV control. Numbers below the western blot images indicate relative pixel densities of the IKKβ or NR4A1 protein levels normalized to corresponding loading controls. **(B)** IKKβ KO AML cells exhibit reduced growth compared to their EV counterpart. Growth curves of EV control, IKKβ pKO1, and pKO2 Monomac-6 cells generated via cell viability measurements using MTS reagent. **(C)** Bar diagram showing the percentage cell cycle distribution of EV, pKO1, and pKO2 MONO-MAC-6 cells measured via BrdU proliferation assay after 2 h treatment of cells with 10 µM BrdU. **(D)** EV, pKO1, and pKO2 MONO-MAC-6 cells were treated with DMSO or 1 µM A9 for 48 h, and viablities of A9-treated cells relative to the corresponding DMSO-treated conditions were assessed. **(E)** EV and NR4A1 KO MOLM-13 cells (described in **Fig. 1P**) were treated with DMSO or 250 nM A9 for 48 h, and viablities of A9-treated cells relative to the corresponding DMSO-treated conditions were assessed. **(F–G)** Principle component analysis (PCA) plots of bulk RNA-sequencing data obtained from MOLM-13 cells **(F)** treated with vehicle (DMSO) vs 0.125 A9 µM or from MONO-MAC-6 cells **(G)** treated with vehicle (DMSO) vs 0.125 A9 µM for 72 h (n = 3 for each treatment group). **(H–I)** Volcano plot depicting differentially expressed genes (DEGs) in A9 treated MOLM-13 (upregulated gene no. = 2, downregulated gene no. = 10, **H**) and MONO-MAC-6 (upregulated gene no. = 25, downregulated gene no. = 136, **I**) cells**.** Data are presented as mean +/- SD. Statistical significance determined using two-way ANOVA followed by Fisher’s LSD test for multiple comparisons between specific groups of interest in **B–E**. *P < 0.05, **P < 0.01, ****P < 0.0001. Only significant comparisons are marked in **C**.

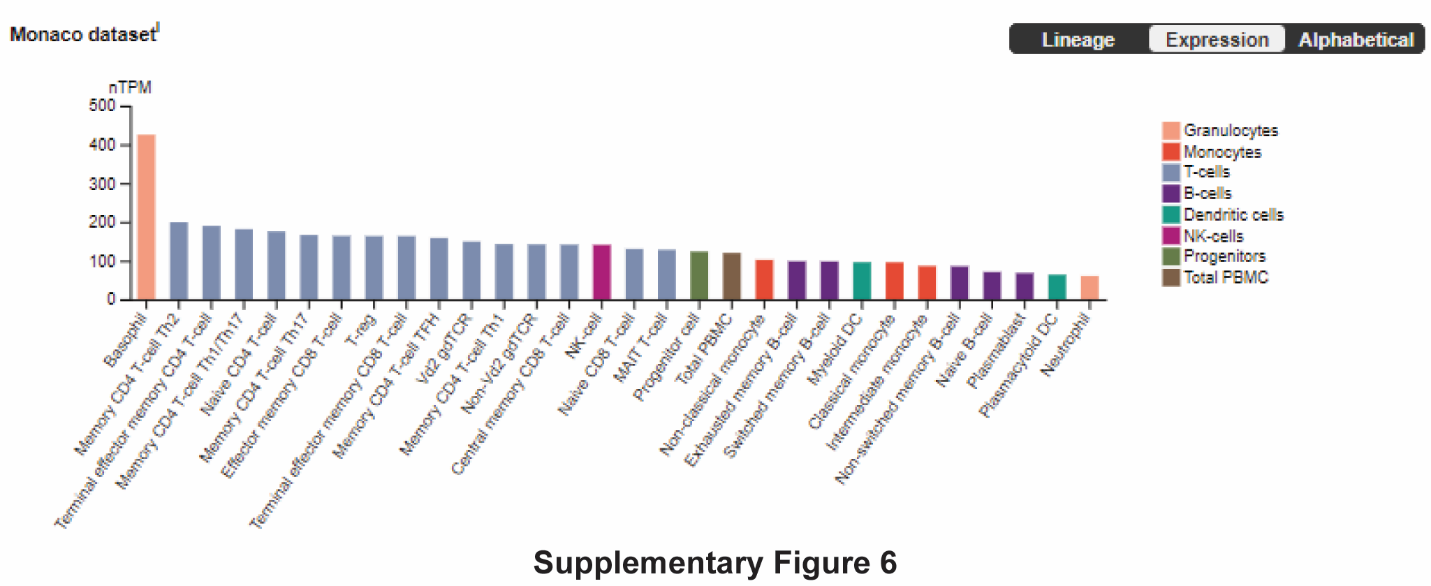

**Supplementary Figure 7:** A dataset from The Human Protein Atlas shows that neutrophils have the least *CRBN* mRNA expression of all different immune cell populations.

**Supplementary Materials and Methods (including detailed methods for the main text)**

**Animals**

7–9-week-old male C57BL6/J mice were purchased from The Jackson Laboratory. Animals were housed in an animal biosafety level 2 (ABSL-2) facility at the University of Florida (UF). All animal handling and experiments were conducted in strict compliance with the UF Institutional Care and Use Committee (IACUC) policies under the protocol number 202400000275. We exercised all efforts to minimize animal suffering.

To evaluate the pathogenicity of EV vs NR4A1 KO human AML cell lines in vivo, 10^6^ MOLM-13 cells (n = 9/group) or 0.5 x 10^6^ MONO-MAC-6 cells (n = 7/group) were injected into the tail vein of male 7–9-week-old male NSG-SGM3 mice (bred inhouse). Blood was withdrawn periodically by puncturing the tail vein and AML cell growth is assessed via flow cytometry using anti-human CD45 (304058/Biolegend) and CD33 (303408/Biolegend) antibodies. End point for euthanasia is determined by symptoms such as drastic weight loss, severe dehydration and anemia, or hind limb paralysis. Mice were euthanized by CO_2_ inhalation.

**KMT2A::MLLT3 mouse AML model**

GFP-tagged KMT2A::MLLT3 were kindly provided by Dr. Chen Zhao at the Case Western Reserve University (CWRU). C57BL6/J mice (n = 4/group) were irradiated with 6.5 Gy X-ray and 24 h later, 2x10^4^ cells were injected per mouse through the tail vein. The first dose of A9 (4 mg/kg) or vehicle was intraperitoneally administered 1 day after cell injections, with subsequent doses administered every 3 days. The A9 dose was reduced to 2 mg/kg after the first four administrations. To evaluate the engraftment and growth of GFP-tagged AML cells in mice, we periodically (twice a week, 30–35 uL per episode) collected blood via tail vein puncture using EDTA treated collection tubes (16.444.10/ Sarttedt). Mouse bloods were immediately mixed with equal volumes of 3 mM EDTA in 1X PBS to prevent coagulation. Further, the blood samples were treated for 10 min with ACK lysis buffer for RBC lysis, washed a couple times with 1X PBS, before running on a flow cytometer to compute GFP+ cell counts per unit volume.

**Complete blood count (CBC) analysis**

Healthy 7-week-old C57BL6/J mice received vehicle (n = 3) or A9 (n = 4) following the regimen described above and were euthanized 19 days after the first injection. For CBC analysis, mouse blood is mixed with 1X PBS containing 2 mM EDTA and run on a HESKA Element HT5 instrument.

**Cells**

MONO-MAC-6, NB-4, and MOLM-13 cells were cultured as suspensions in RPMI-1640 media supplemented with 10% FBS; HL-60 and U-937 in IMDM + 10% FBS; HEK293T and A375 cells in DMEM + 10% FBS. AML cell lines were generously provided by Dr. Zhijian Qian (formerly at UF). HEK293T and A375 cells were acquired from ATCC and cultured in DMEM media supplemented with 10% FBS. All culture media contained 100 units/mL of penicillin and 100 µg/mL streptomycin. Saturated cell cultures (with concentrations greater than 1-2 million cells/mL) were split at 1:10 ratio for maintenance.

KMT2A::MLLT3 cells were cultured in IMDM media containing 10% FBS media, 100 units/mL of penicillin and 100 µg/mL streptomycin and supplemented with 10 ng/mL murine interleukin-3 (IL-3), IL-6, granulocyte-macrophage colony-stimulating factor (GM-CSF), and stem cell factor (SCF). P401 cells (public model name AML001) were from an AML patient (IRB H-3342) characterized by a chromosomal insertion: ins(10;11) [2]. DFAM6855 was purchased from PRoXe (public repository of xenografts) and is a therapy-related AML (IRB 20210216) with a complex karyotype including t(9;11)(KMT2A::MLLT3). DFAM68555 and P401, were treated with 250 nM celastrol, A9 NC, or A9 in RPMI+10% FBS before counting live cells via trypan blue exclusion method.

**Supplementary Table 1:** List of human AML cell lines used and their brief information.

| **Human AML Cell Line** | **Characteristics** |
| --- | --- |
| NB-4 | Acute promyelocytic leukemia cell line cytogenetically characterized by a t(15;17) (q22;q11-12) translocation [3] |
| MONO-MAC-6 | Monoblastic leukemia cell line with phenotypic and functional features of mature monocytes [4] |
| MOLM-13 | Acute monocytic leukemia cell line that express monocyte-specific esterase (MSE) and *MLL-AF9* fusion mRNA [5] |
| U-937 | Promonocytic human myeloid leukemia cell line isolated from histiocytic lymphoma of a 37-year-old male [6] |
| HL-60 | Continuous, differentiating myeloid cell line from a patient with acute promyelocytic leukemia [7] |

**Chemistry**

The chemical structures, detailed synthetic procedures, and nuclear magnetic resonance spectra of PROTACs are presented in supplementary information.

**MTS assay**

AML cells were plated onto 96-well plates (at 10000–20000 cells/well) and treated with drugs or DMSO. 20 µL of 1:20 PMS (AC130160010/Fisher scientific) and MTS (G1111/Promega) mixture was added to 200 µL cell suspension/well. After 4 h incubation, absorbances were recorded at 490 nm using Biotek Synergy Neo2 Multimode Reader.

**PE-Annexin/7-AAD labeling**

Annexin/7-AAD labeling was performed according to the manufacturer (88-8102-72/eBiosciences) instructions with some modifications on the concentration of reagents. Briefly, cells were washed in 1X binding buffer, stained with PE-conjugated Annexin V (0.5 µL/100 µL buffer) at room temperature for 10–15 min, washed again in 1X binding buffer before finally staining with 7-AAD Viability Staining Solution (5 µL/ 200 µL buffer). The samples were immediately run on a flow cytometer for analysis.

**BrdU proliferation assay**

BrdU/7-AAD labeling was performed by exactly following the manufacturer instructions (BDB559619/BD Biosciences). Briefly, BrdU was added to cell culture at 10 µM final concentration and incubated for 2 h for pulse-labeling. The cells were then subsequently washed, fixed and permeabilized, treated with DNase to expose incorporated BrdU, stained with FITC-conjugated anti-BrdU antibody and finally labeled with 7-AAD before running on a flow cytometer.

**Knockdown studies**

siRNAs were used for *IKBKB* and *NR4A1* knockdown in NB-4 cells. ON-TARGETplus siRNAs targeting h*NR4A1* (L-003426-00-0005) and h*IKBKB* (L-003503-00-0005) as well as the non-targeting control (D-001910-01-05) were ordered from Horizon Discovery. NB-4 cells were nucleofected with 20 nM total siRNA using SF cell line 4D nucleofector X kit (EH-100 pulse code, Lonza) using the manufacturer instructions.

However, for knocking down *IKBKB* and *NR4A1* knockdown in MONO-MAC-6 and MOLM-13 cells, predesigned shRNA lentiviral vectors (TRCN0000019427/Sigma-Aldrich for *NR4A1* and TRCN0000018917/Sigma-Aldrich for *IKBKB*) were used. Scrambled shRNA pLKO.1 vector (1864/Addgene) was used as an empty vector (EV) control. For lentiviral production, EV or shRNA lentiviral vector, pSPAX2 plasmid (12260/Addgene) and pVSVg (8454/Addgene) were transfected into HEK293T cells. 48 h later, supernatants from the transfected cultures were collected, filtered through 0.45 µm strainer, and combined with AML (Monomac-6 and Molm-13) cell suspensions in 2:1 ratio to facilitate lentiviral transduction. The cells were centrifuged and cultured in fresh media after 48 h. Knockdown was verified after additional 48 h via qRT-PCR or western blotting.

**qRT-PCR**

RNA was prepared from AML cells using a commercial RNA isolation kit (740955.5/Macherey-Nagel). cDNA was synthesized from 100–200 ng RNA using SuperScript^®^ III First Strand cDNA preparation kit (Invitrogen). Diluted cDNA was then used for qPCR (cycling conditions: denaturation at 95 °C for 10 min followed by 40 cycles of 95 °C for 15 s and 60 °C for 1 min) with gene-specific primers in the presence of PowerUp™ SYBR™ Green Master Mix for qPCR (A25742/Thermofisher) on an Applied Biosystems QuantStudio 3 instrument. Fold changes in each gene mRNA levels were calibrated to *GAPDH* mRNA expression and computed using the 2^-∆∆Ct^ method.

The following primers were used in qRT-PCR:

*NR4A1*

Forward: 5’-AGGGCTGCAAGGGCTTCT-3’

Reverse: 5’-GGCAGATGTACTTGGCGTTTTT-3’

*IKBKB*

Forward: 5'-CCGACAGAGTTAGCACGACA-3'

Reverse: 5'-GGCAATCTGCTCACCTGTTT-3'

*gapdh*

Forward: 5'-CTGGGCTACACTGAGCACC-3'

Reverse: 5'-AAGTGGTCGTTGAGGGCAATG-3'

**Western blotting**

Suspension cultures of AML cells were harvested and lysed with cold RIPA buffer (50 mM Tris, pH 8.0, 150 mM NaCl, 1% Triton X-100, 0.1% SDS, 0.5% sodium deoxycholate containing protease and phosphatase inhibitor cocktail (PPC1010/Sigma). Protein concentrations of the cell extracts were determined by using Pierce™ BCA Protein Assay (23227/Thermo Scientific). Lysate volumes were adjusted to obtain the same final concentrations. Equal amounts of protein per sample were separated by SDS-PAGE and transferred onto nitrocellulose membranes (1620115/BioRad). Membranes were blocked with 5% non-fat milk in TBS-T (Tris-buffered saline containing 0.05% Tween-20) and incubated overnight at 4 °C on a shaker with primary antibody solutions prepared with 3% BSA (bovine serum albumin) in TBS-T. After three 10 min washes, the membranes were incubated with suitable secondary antibodies prepared in 5% milk in TBS-T. Proteins were detected using SuperSignal™ West Pico PLUS Chemiluminescent Substrate (34580/Thermo Scientific).Primary antibodies: anti-IKKβ (2684/Cell Signaling, 1:1000 dilution); anti-NR4A1/Nur77 (ab283264/Abcam, 1:1000); anti-β-tubulin (2146/Cell Signaling, 1:5000); anti-GAPDH (5174/Cell Signaling, 1:5000); anti-CRBN (71810/Cell Signaling, 1:1000); anti-caspase 3 (ab4051/Abcam, 1:1000); anti-cleaved caspase 3 (9661/Cell Signaling, 1:1000), anti-PI3Kα (4249/Cell Signaling, 1:1000); anti-Chk1 (2360/Cell Signaling, 1:1000); M2 anti-flag (F1804/Sigma, 1:1000) and anti-RIP (4926/Cell Signaling, 1:1000). Secondary: HRP anti-rabbit IgG (7074/Cell Signaling, 1:2000) and HRP anti-mouse IgG (7076/Cell Signaling, 1:2000).

**Co-immunoprecipitation**

For A9-induced co-IP of IKKβ and NR4A1 with CRBN, MONO-MAC-6 cells were initially treated for 1 h with 5 µM MG132 followed by co-treatment of 10 µM A9 or A9 NC for additional 3 h. Protein extracts (550 µg total protein) in NP-40 buffer containing protease and phosphatase inhibitor cocktail (PPC1010/Sigma) were incubated overnight with 30 µL protein A Sepharose beads, and 4 µg/mL anti-CRBN antibody (71810/Cell Signaling) overnight. The beads were washed and boiled with 1x SDS PAGE buffer for western blotting.

For A9-induced CRBN-IKKβ co-immunoprecipitation, HEK293T cells were transfected with pMAX-GFP (177825/Addgene) or Flag-CRBN. Flag-CRBN expressing cells were treated with 5 μM MG132 for 1 h before adding celastrol, A9, or A9 NC at 10 µM concentration for further 3 h incubation. Protein extracts in NP-40 buffer were incubated with anti-Flag magnetic beads (M8823/Sigma) overnight at 4 °C under rotating motion. The beads were washed with NP-40 buffer and immunoprecipitated proteins were separated by SDS-PAGE. The membranes were immunoblotted for CRBN, IKKβ, or GAPDH.

For A9-induced NR4A1-CRBN co-immunoprecipitation, Flag-NR4A1 (HG17699-CF/SinoBiological) was expressed in HEK293T cells followed by the above procedure. Finally, the membranes were immunoblotted for CRBN, Flag, or GAPDH.

**Flow cytometry**

NB-4 and MONO-MAC-6 cells were plated on 6-well plates at 5 x 10^5^ cells in 2 mL RPMI complete media per well and treated with DMSO or A9 at defined concentrations and durations (as indicated in the corresponding figure legends). Following treatment, the cells were collected in 15 mL conical tubes and washed a couple times with 1X PBS before labeling with Live/Dead Zombie Red viability dye (423110/Biolegend, 1:1000) and BV510 anti-human CD45 (304036/Biolegend, 1:100) in PBS. Further, the cells were fixed (using a 1:3 mixture of Fixation/Permeabilization Concentrate, 00-5123-43/Thermofisher and the Fixation/Permeabilization Diluent, 00-522/Thermofisher); permeabilized (using Permeabilization Buffer, 00-8333/Thermofisher, diluted to 1X); and stained with PE anti-NR4A1 antibody (59999/Cell Signaling, 1:50) or PE anti-IKKβ antibody (NB100-56509PE/Novus). After final washes with FACS buffer (2% FBS in PBS), the cells were run on a Cytek Aurora flow cytometer. Data analysis was performed in FlowJo software to evaluate the NR4A1 protein levels in live AML cells with or without A9 treatment.

For cell differentiation assays, MOLM-13 and MONO-MAC-6 cells were washed in 1x PBS and stained at 4 °C for 30 min with Live/Dead Zombie Red viability dye (423110/Biolegend, 1:1000), APC/Cy7 anti-CD14 (325619/Biolegend, 3:100), PerCP/Cy5.5 anti-CD15 (323020/Biolegend, 3:100) and PE/Dazzle 594 anti-CD11b (101256, Biolegend, 2:100) in the presence of human FcR blocker (130-059-901/Miltenyi, 1:100). The cells were washed with FACS buffer before running on a cytometer.

***IKBKB* and *NR4A1* knockout**

Single guide RNA (sgRNA) targeting *IKBKB* or *NR4A1* (+overhangs, shown below) were cloned into lentiCRISPRv2 plasmid (52961, Addgene) as previously described [8, 9]. Guide oligos targeting the human *IKBKB* gene were designed using the Synthego guide designing tool.

*IKBKB* guides

Oligo1 forward: CACCGACATTGGGGTGGGTCAGCCT

Oligo1 reverse: AAACAGGCTGACCCACCCCAATGTC

Oligo2 forward: CACCGGGCAGCCACCACATTGGGGT

Oligo2 reverse: AAACACCCCAATGTGGTGGCTGCCC

*NR4A1* guide

Oligo1 forward: CACCGCTGGTGTCCCATATTGGGCT

Oligo1 reverse: AAACAGCCCAATATGGGACACCAGC

For lentiviral production, lentiCRISPRv2 plasmid with or without sgRNA insert, pSPAX2 plasmid (12260/Addgene) and pVSVg (8454/Addgene) were transfected into HEK293T cells. 48 h later, supernatants from the transfected cultures were collected, filtered through 0.45 µm strainer, and combined with AML (Monomac-6 and Molm-13) cell suspensions in 1:1 ratio to facilitate lentiviral transduction. Transduced cells were initially selected with 0.4 µg/mL puromycin, which was further reduced to 0.2 µg/mL for the maintenance of pooled KO cells.

**Patient Cohorts**

The analyzed cohorts were: 153 patients from the TCGA-LAML project for whom gene expression and survival data was available from the NCI GDC portal (https://portal.gdc.cancer.gov/); 163 patients from the St. Jude AML02 trial (NCT00136084) and 1,048 from the TARGET-AML initiative for whom gene expression and survival data was available. The study design and outcomes for the pediatric AML trial St. Jude AML02 (NCT00136084) have been previously reported [10]. TARGET-AML contains data from Children’s Oncology Group studies and is available at: <https://portal.gdc.cancer.gov/projects/TARGET-AML>.

**Bioinformatic analysis**

UMAP plots were derived from the Single Cell Portal (<https://singlecell.broadinstitute.org/single_cell/study/SCP1987>) dataset deposited by Lasry et al [1]. The Kaplan-Meier method was used to estimate the OS probabilities for patients in the analysis cohorts based on NR4A1 and IKBKB expression groups by median or quartiles. Cox proportional hazard models were used for association with OS. All survival association analyses were performed using R version 4.3.1 (R Project for Statistical Computing). Proteomics were performed and analyzed at the IDeA National Resource for Quantitative Proteomics, University of Arkansas for Medical Sciences.

**Single-cell RNA sequencing analysis**

scRNA-seq datasets derived from seven healthy controls and ten AML patients (GSE185381) were subjected to standard scRNA-seq data processing, quality control, integration, and bioinformatic analysis using R tookit Seurat (v.4.0.6) as previously described [11]. Gene set enrichment analysis (GSEA) was conducted with the escape R package (v.1.6.0) [12]. Correlation analysis was performed using the FeatureScatter function on Seurat. The cor.test() function was used to calculate the statistical significance of the correlation.

**Bulk RNA sequencing analysis**

For RNAseq, RNA from DMSO and A9 treated cells (n = 3) was purified and sent to the University of Florida Interdisciplinary Center for Biotechnology Research Gene Expression and Genotyping Core (RRID: SCR_019145). RNA concentration was determined by Qubit and quality was determined using a Bioanalyzer RNA assay. Samples with a RIN score ≥ 8 were used for library preparation. Library preparation was done using the NEBNext Poly(A) mRNA isolation module (New England Biolabs) and NEBNext Ultra II Library prep Kit (New England Biolabs). Libraries were sequenced at 150 paired end reads at 25 million reads/sample using an Illumina Novaseq 6000 (Illumina). Raw FastQ sequencing data were aligned to the hg38 genome using *Kallisto* [13]. Differential gene expression analysis was conducted with the *limma* package [14], and pathway enrichment was evaluated using complete ranked gene set in GSEA software [15]. *ggplot2* package in R was used for data visualization and graph generation.

**Proteomics**

MONO-MAC-6 cells were treated in triplicate with DMSO, 0.5 µM celastrol, or A9 for 16 h before for cell lysate preparation using cold RIPA buffer (50 mM Tris, pH 8.0, 150 mM NaCl, 1% Triton X-100, 0.1% SDS, 0.5% sodium deoxycholate containing protease and phosphatase inhibitor cocktail (PPC1010/Sigma). Protein concentrations were measured and 80 µg protein (at least 30 required) per sample were sent to IDeA National Resource for Quantitative Proteomics at University of Arkansas for Medical Sciences.

**Statistics**

Sample size determination did not involve any statistical methods. 3 biological replicates per group were generally used for in vitro studies, while the number of animals per group varied from 3–9 depending on the study with exact numbers reported in the figure legends as well as in the ‘Animals’ section. For in vivo studies, animals were randomly assigned into experimental groups. Investigators were not blinded to allocation during experiments and outcome assessments. No animals were excluded from the analysis. Statistical tests were performed in GraphPad Prism Version 10.6.0 (890). Unpaired two-tailed T-test was used compare a variable between two groups, while one-way ANOVA with Tukey’s test to correct for multiple comparisons used to compare more than two groups. Correction for multiple comparison was skipped in cases when the comparisons of interest were predetermined. Studies with multiple variables and groups were analyzed using two-way ANOVA. Non-linear regression was used to generate dose response curves and compute IC50 values. Log-rank test was used for survival analysis in animal experiments.

**Mutant IKKβ expression in MONO-MAC-6**

Lentiviral vector to express lysine-mutant IKKβ (with sequence below) was custom designed by Twist Bioscience. Briefly, insert DNA was cloned into a pTwist Lenti CMV Puro. Mutant IKKβ expression vector and a lentiviral control (1864/Addgene) vector were separately packaged into lentivirus, which was used to transduce MONO-MAC-6 cells as described in detail above (in the knockdown studies section). Transduced cells were selected under 1 µg/mL puromycin after 5 days and used for MTS assays.

**Protein sequence of lysine-mutant IKKβ**

```
IKK2-K147R,301R, 418R, 555R, 703R

MSWSPSLTTQTCGAWEMKERLGTGGFGNVIRWHNQETGEQIAIK 44
                     QCRQELSPRNRERWCLEIQIMRRLTHPNVVAARDVPEGMQNLAPNDLPLLAMEYCQGG 58 102
                     DLRKYLNQFENCCGLREGAILTLLSDIASALRYLHENRIIHRDLRPENIVLQQGEQRL 58 160 K147R
                     IHKIIDLGYAKELDQGSLCTSFVGTLQYLAPELLEQQKYTVTVDYWSFGTLAFECITG 58 218
                     FRPFLPNWQPVQWHSKVRQKSEVDIVVSEDLNGTVKFSSSLPYPNNLNSVLAERLEKW 58 276
                     LQLMLMWHPRQRGTDPTYGPNGCFRALDDILNLKLVHILNMVTGTIHTYPVTEDESLQ 58 334 K301R
                     SLKARIQQDTGIPEEDQELLQEAGLALIPDKPATQCISDGKLNEGHTLDMDLVFLFDN 58 392
                     SKITYETQISPRPQPESVSCILQEPRRNLAFFQLRKVWGQVWHSIQTLKEDCNRLQQG 58 450 K418R
                     QRAAMMNLLRNNSCLSKMKNSMASMSQQLKAKLDFFKTSIQIDLEKYSEQTEFGITSD 58 508
                     KLLLAWREMEQAVELCGRENEVKLLVERMMALQTDIVDLQRSPMGRRQGGTLDDLEEQ 58 566 K555R
                     ARELYRRLREKPRDQRTEGDSQEMVRLLLQAIQSFEKKVRVIYTQLSKTVVCKQKALE 58 624
                     LLPKVEEVVSLMNEDEKTVVRLQEKRQKELWNLLKIACSKVRGPVSGSPDSMNASRLS 58 682
                     QPGQLMSQPSTASNSLPEPARKSEELVAEAHNLCTLLENAIQDTVREQDQSFTALDWS 58 740 K703R

WLQTEEEEHSCLEQAS
```

**Chemical Synthesis**

Dimethylformamide (DMF) and dichloromethane (DCM) were obtained via a solvent purification system by filtering through two columns packed with activated alumina and 4 Å molecular sieve, respectively. Water was purified with a Milli-Q Simplicity 185 Water Purification System (Merck Millipore). All other chemicals and solvents obtained from commercial suppliers were used without further purification. Reaction progress was monitored by thin-layer chromatography (silica-coated glass plates) and visualized by 256 nm and 365 nm UV light, and/or by liquid chromatography–mass spectrometry (LC-MS). Flash chromatography was performed using silica gel (230-400 mesh) as the stationary phase on the Biotage Isolera purification system. Preparative thin-layer chromatography (Prep-TLC) was performed on silica gel 60 F254 20 cm x 20 cm glass plates from Sigma-Aldrich. ^1^H nuclear magnetic resonance (NMR) spectra were recorded in CDCl_3_ or CD_3_OD at 600 MHz, and ^13^C-NMR spectra were recorded at 151 MHz using a Bruker (Billerica, MA) DRX NMR spectrometer. Chemical shifts δ are given in ppm using tetramethylsilane as an internal standard. Multiplicities of NMR signals are designated as singlet (s), doublet (d), doublet of doublets (dd), triplet (t), quartet (q), multiplet (m), and broad singlet (bs). All final compounds for biological testing were of 95.0% purity as analyzed by LC-MS, performed on an Advion AVANT LC system with the expression Compact Mass Spectrometer (CMS) using a Thermo AccucoreTM VanquishTM C18+ UHPLC Column (1.5 µm, 50 x 2.1 mm) at 40 °C. Gradient elution was used for UHPLC with a mobile phase of acetonitrile and water containing 0.1% formic acid.
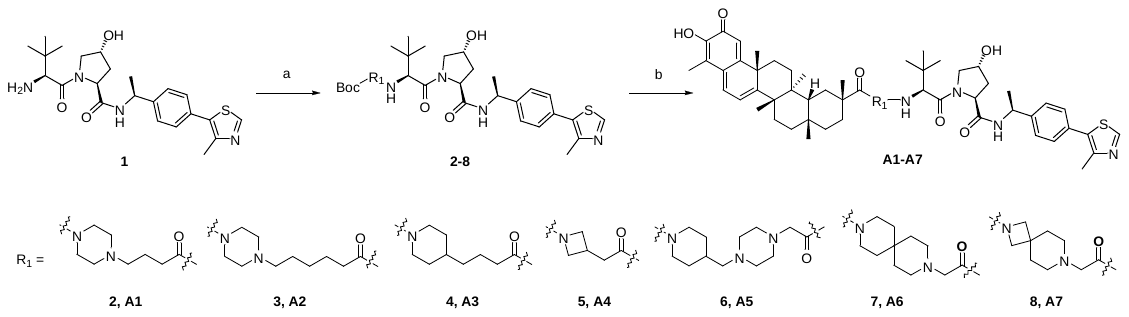

Scheme 1. Synthesis of **A1-A7**. Reagents and conditions: (a) HATU, DIPEA, DCM, rt, overnight; (b) i) TFA, DCM, ii) Celastrol, HATU, DIPEA, DMF, rt, overnight.

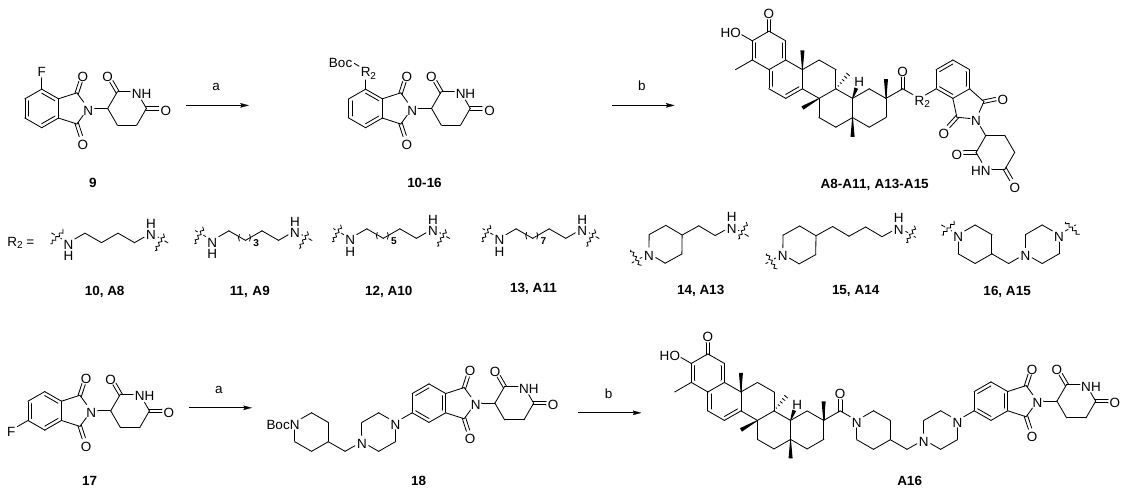

Scheme 2. Synthesis of **A8-A11**, **A13-A16**. Reagents and conditions: (a) DIPEA, DMSO, 90 °C, overnight; (b) i) TFA, DCM, ii) Celastrol, HATU, DIPEA, DMF, rt, overnight.

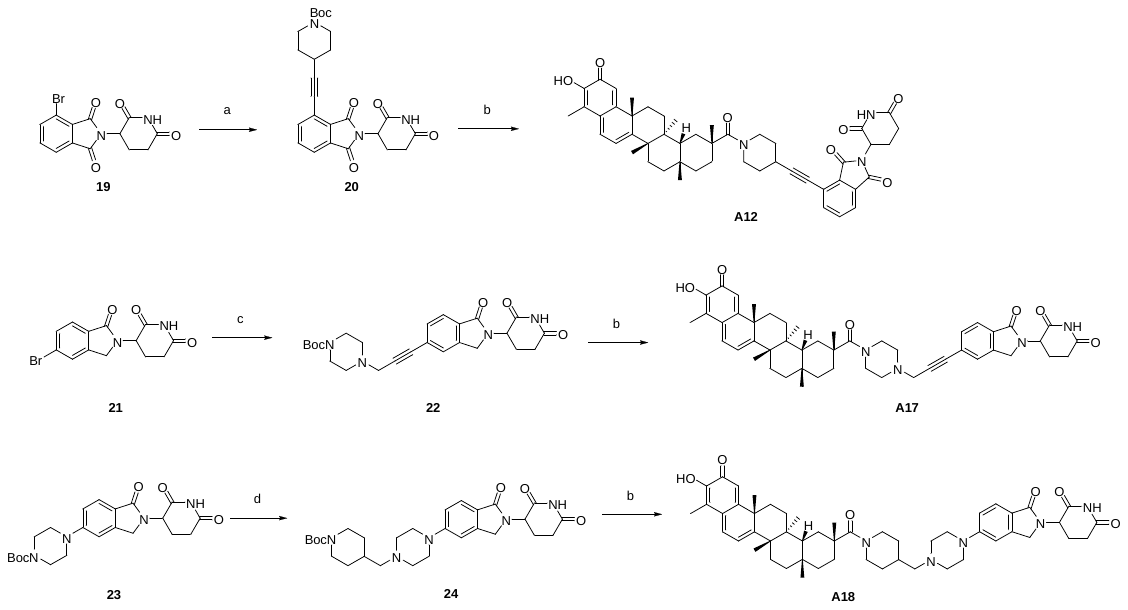

Scheme 3. Synthesis of **A12, A17-A18**. Reagents and conditions: (a) tert-butyl 4-ethynylpiperidine-1-carboxylate, Pd(PPh_3_)_2_Cl_2_,CuI, Et_3_N, DMF, 80 °C, overnight; (b) i) TFA, DCM, ii) Celastrol, HATU, DIPEA, DMF, rt, overnight; (c) tert-butyl 4-(prop-2-yn-1-yl)piperazine-1-carboxylate, Pd(PPh_3_)_2_ Cl_2_,CuI, Et_3_N, DMF, 80 °C; (d) i) TFA, DCM, ii) tert-butyl 4-formylpiperidine-1-carboxylate, NaBH(OAc)_3_, Et_3_N, DCM, rt, overnight

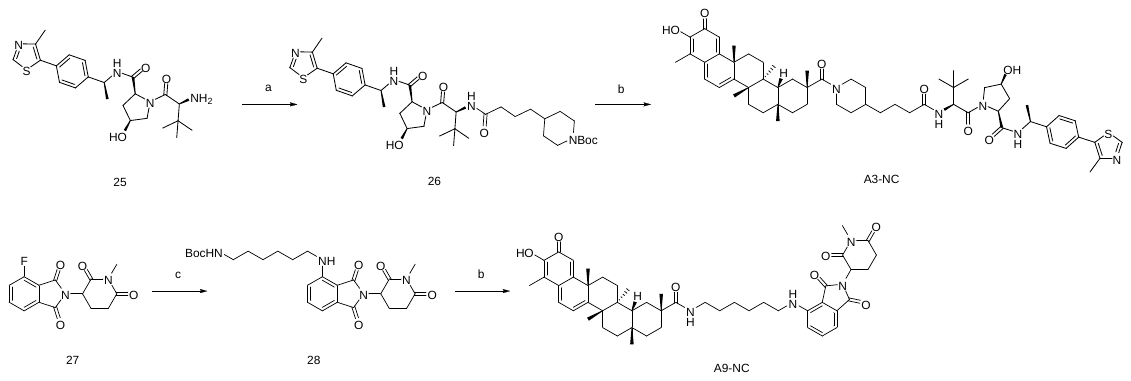

Scheme 4. Synthesis of **A3-NC**, **A9-NC**. Reagents and conditions: (a) HATU, DIPEA, DCM, rt, overnight; (b) i) TFA, DCM, ii) Celastrol, HATU, DIPEA, DMF, rt, overnight; (c) DIPEA, DMSO, 90 °C, overnight.

**General Procedure A (for the preparation 2-8).**

To a solution of VHL-amine HCl salt (1.0 equiv) and corresponding acid-terminated linkers (1.0 equiv) in 5 mL DCM was added hexafluorophosphate azabenzotriazole tetramethyl uranium (HATU) (1.1 equiv) and N, N-diisopropylethylamine (DIPEA) (5.0 equiv), and the reaction was stirred at room temperature overnight. The reaction mixture was poured into a saturated NH_4_Cl solution, and the aqueous layer was extracted with ethyl acetate. The combined organic layers were washed with brine, dried over Na_2_SO_4_, and concentrated. The crude product was puriﬁed by silica gel column chromatography to afford the products.

**General Procedure B (for the preparation 10-16, 18).**

To a solution of 9 or 17 (1.0 equiv) in DMSO (3 mL) was added corresponding amine-terminated linkers (1.0 equiv) and DIPEA (5 equiv). The mixture was stirred at 90 °C overnight. The mixture was diluted with 20 mL of ethyl acetate, and the solution was washed with water (20 mL × 3). The organic layers were washed with brine (20 mL), dried over Na_2_SO_4_, filtered, and concentrated in vacuo. The crude product was puriﬁed by silica gel column chromatography to aﬀord the products.

**General Procedure C (for the preparation A1-A18, A3-NC, A9-NC).**

Triﬂuoroacetic acid (0.5 mL) was added to a solution of tert-butyl ester compound (2-8, 10-16, 18, 20, 22, 24, 26, 28, 0.034 mmol, 1.0 equiv) in DCM (1 mL). Then, the mixture was stirred at room temperature for 1 h. After TLC monitoring, the mixture was concentrated in vacuo and directly used in the next step without further puriﬁcation. The above crude was dissolved in 1 mL of DMF, and celastrol was added followed by DIPEA (0.17 mmol, 5.0 equiv) and HATU (0.037 mmol, 1.1 equiv). The reaction mixture was then stirred at room temperature overnight. The mixture was diluted with 20 mL of water, and the solution was extracted with ethyl acetate (10 mL × 3). The combined organic layers were washed with brine (10 mL × 3), dried over Na_2_SO_4_, filtered, and concentrated in vacuo. The crude product was puriﬁed by silica gel column chromatography to aﬀord the products.

Intermediates **2** to **5** were prepared as reported [16]

**tert-Butyl 4-((4-(2-(((S)-1-((2S,4R)-4-hydroxy-2-(((S)-1-(4-(4-methylthiazol-5-yl)phenyl)ethyl)carbamoyl)pyrrolidin-1-yl)-3,3-dimethyl-1-oxobutan-2-yl)amino)-2-oxoethyl)piperazin-1-yl)methyl)piperidine-1-carboxylate (6)**

Following the general procedure A, compound **6** (120 mg, 71%) was obtained as white solid. ^1^H NMR (600 MHz, CDCl_3_) δ 8.68 (s, 1H), 7.89 (d, *J* = 8.2 Hz, 1H), 7.46 (d, *J* = 7.8 Hz, 1H), 7.43 – 7.33 (m, 4H), 5.08 (p, *J* = 7.1 Hz, 1H), 4.76 (t, *J* = 7.9 Hz, 1H), 4.50 (s, 1H), 4.42 (d, *J* = 8.2 Hz, 1H), 4.21 (dt, *J* = 11.4, 1.9 Hz, 1H), 4.08 (s, 2H), 3.58 (dd, *J* = 11.5, 3.6 Hz, 1H), 3.01 (q, *J* = 16.6 Hz, 2H), 2.69 – 2.66 (m, 2H), 2.60 – 2.39 (m, 11H), 2.19 (d, *J* = 7.1 Hz, 2H), 2.09 – 2.05 (m, 1H), 1.75 – 1.60 (m, 7H), 1.45 (s, 12H), 1.07 (s, 9H). ESI [M + H]+ (m/z): 768.5.

**tert-Butyl 9-(2-(((S)-1-((2S,4R)-4-hydroxy-2-(((S)-1-(4-(4-methylthiazol-5-yl)phenyl)ethyl)carbamoyl)pyrrolidin-1-yl)-3,3-dimethyl-1-oxobutan-2-yl)amino)-2-oxoethyl)-3,9-diazaspiro[5.5]undecane-3-carboxylate (7)**

Following the general procedure A, compound **7** (90 mg, 60%) was obtained as white solid.

^1^H NMR (600 MHz, CDCl_3_) δ 8.66 (s, 1H), 7.89 (d, *J* = 8.6 Hz, 1H), 7.47 (d, *J* = 7.4 Hz, 1H), 7.40 – 7.31 (m, 4H), 5.06 (t, *J* = 7.1 Hz, 1H), 4.72 (t, *J* = 7.9 Hz, 1H), 4.56 – 4.37 (m, 2H), 4.09 (dd, *J* = 11.1, 7.3 Hz, 1H), 3.60 (dd, *J* = 11.3, 3.8 Hz, 1H), 3.39 – 3.31 (m, 4H), 3.04 – 2.91 (m, 2H), 2.52 – 2.41 (m, 8H), 2.09 – 2.01 (m, 1H), 1.57 – 1.33 (m, 21H), 1.04 (s, 9H). ESI [M + H]^+^ (*m*/*z*): 739.5.

**tert-Butyl 7-(2-(((S)-1-((2S,4R)-4-hydroxy-2-(((S)-1-(4-(4-methylthiazol-5-yl)phenyl)ethyl)carbamoyl)pyrrolidin-1-yl)-3,3-dimethyl-1-oxobutan-2-yl)amino)-2-oxoethyl)-2,7-diazaspiro[3.5]nonane-2-carboxylate (8)**

Following the general procedure A, compound **8** (110 mg, 64%) was obtained as white solid. ^1^H NMR (600 MHz, CDCl_3_) δ 8.68 (s, 1H), 7.85 (d, *J* = 8.5 Hz, 1H), 7.45 (d, *J* = 7.7 Hz, 1H), 7.43 – 7.34 (m, 4H), 5.10 - 5.05 (m, 1H), 4.75 - 4.72 (m, 1H), 4.51 - 4.47 (m, 2H), 4.12 – 4.05 (m, 1H), 3.73 - 3.70 (m, 1H), 3.65 – 3.60 (m, 4H), 3.03 – 2.90 (m, 2H), 2.56 – 2.38 (m, 9H), 2.09 – 2.02 (m, 1H), 1.79 (s, 4H), 1.50 – 1.40 (m, 12H), 1.06 (s, 9H). ESI [M + H]^+^ (*m*/*z*): 711.5.

Intermediates **10** to **13, 16, 18** were prepared as reported [17-19]

**tert-Butyl 4-(2-((2-(2,6-dioxopiperidin-3-yl)-1,3-dioxoisoindolin-4-yl)amino)ethyl)piperidine-1-carboxylate (14).**

Following the general procedure B, compound **14** (180 mg, 61%) was obtained as yellow solid.

^1^H NMR (600 MHz, CDCl3) δ 7.51 – 7.48 (m, 1H), 7.11 (d, *J* = 7.2 Hz, 1H), 6.88 (d, *J* = 8.5 Hz, 1H), 6.19 (t, *J* = 5.5 Hz, 1H), 4.10 (s, 1H), 3.35 - 3.28 (m, 2H), 2.96 – 2.65 (m, 6H), 1.83 - 1.65 (m, 4H), 1.65 - 1.52 (m, 4H), 1.46 (s, 9H), 1.18 (d, *J* = 7.2 Hz, 2H). ESI [M + H]^+^ (*m*/*z*): 485.5.

**tert-Butyl 4-(4-((2-(2,6-dioxopiperidin-3-yl)-1,3-dioxoisoindolin-4-yl)amino)butyl)piperidine-1-carboxylate (15)**

Following the general procedure B, compound **15** (200 mg, 65%) was obtained as yellow solid.

^1^H NMR (600 MHz, CDCl_3_) δ 7.50 – 7.47 (m, 1H), 7.09 (dd, *J* = 7.6, 2.2 Hz, 1H), 6.88 (d, *J* = 8.5 Hz, 1H), 4.96 – 4.90 (m, 1H), 4.06 (s, 2H), 3.47 – 3.39 (m, 2H), 3.27 (td, *J* = 7.0, 5.3 Hz, 1H), 2.93 – 2.55 (m, 6H), 2.18 – 2.09 (m, 1H), 1.66 (p, *J* = 7.3 Hz, 2H), 1.60 – 1.52 (m, 2H), 1.45 (s, 9H), 1.32 – 1.19 (m, 4H), 1.13 – 0.98 (m, 2H). ESI [M + H]^+^ (*m*/*z*): 513.5.

**tert-Butyl 4-((2-(2,6-dioxopiperidin-3-yl)-1,3-dioxoisoindolin-4-yl)ethynyl)piperidine-1-carboxylate (20)**

To a solution of **19** (135 mg, 0.4 mmol) in 3 mL of DMF was added CuI (15 mg, 0.08 mmol) and Et_3_N (121 mg, 1.2 mmol). Then, tert-butyl 4-ethynylpiperidine-1-carboxylate (84 mg, 0.4 mmol) and Pd (PPh_3_) _2_Cl_2_ (29 mg, 0.04 mmol) were added into the mixture under Ar protection. The mixture was stirred at 80 °C under Ar atmosphere overnight. The reaction mixture was poured into a saturated NH_4_Cl solution, and the aqueous layer was extracted with ethyl acetate. The combined organic layers were washed with brine, dried over Na_2_SO_4_, and concentrated. The crude product was puriﬁed by ﬂash column chromatography to aﬀord the compound **20** (89 mg, 75 % yield) as a white solid. ^1^H NMR (600 MHz, CDCl_3_) δ 7.99 (s, 1H), 7.79 (dd, *J* = 7.2, 1.2 Hz, 1H), 7.73 – 7.64 (m, 2H), 4.99 (dd, *J* = 12.6, 5.4 Hz, 1H), 3.79 – 3.72 (m, 2H), 3.38 – 3.31 (m, 2H), 2.96 – 2.70 (m, 4H), 2.20 – 2.11 (m, 1H), 1.89 (d, *J* = 3.9 Hz, 2H), 1.80 – 1.71 (m, 2H), 1.47 (s, 9H). ESI [M + H]^+^ (*m*/*z*): 366.4.

**tert-Butyl 4-(3-(2-(2,6-dioxopiperidin-3-yl)-1,3-dioxoisoindolin-5-yl)prop-2-yn-1-yl)piperazine-1-carboxylate (22)**

To a solution of **21** (103 mg, 0.32 mmol) in 3 mL of DMF was added CuI (12 mg, 0.064 mmol) and Et_3_N (97 mg, 0.96 mmol). Then, tert-butyl 4-(prop-2-yn-1-yl)piperazine-1-carboxylate (72 mg, 0.32 mmol) and Pd (PPh_3_) _2_Cl_2_ (23 mg, 0.032 mmol) were added into the mixture under Ar protection. The mixture was stirred at 80 °C under Ar atmosphere overnight. The reaction mixture was poured into a saturated NH_4_Cl solution, and the aqueous layer was extracted with ethyl acetate. The combined organic layers were washed with brine, dried over Na_2_SO_4_, and concentrated. The crude product was puriﬁed by ﬂash column chromatography to aﬀord the compound **22** (89 mg, 75 % yield) as a white solid. ^1^H NMR (600 MHz, CDCl_3_) δ 8.06 (s, 1H), 7.82 (d, *J* = 7.9 Hz, 1H), 7.57 – 7.50 (m, 2H), 5.21 (dd, *J* = 13.4, 5.1 Hz, 1H), 4.48 (d, *J* = 16.0 Hz, 1H), 4.32 (d, *J* = 16.0 Hz, 1H), 3.56 (s, 2H), 3.51 (s, 4H), 2.99 – 2.78 (m, 2H), 2.59 (s, 4H), 2.40 - 2.32 (m, 1H), 2.25 - 2.21 (m, 1H), 1.47 (s, 9H). ESI [M + H]^+^ (*m*/*z*): 467.2.

**tert-Butyl 4-((4-(2-(2,6-dioxopiperidin-3-yl)-1-oxoisoindolin-5-yl)piperazin-1-yl)methyl)piperidine-1-carboxylate (24)**

To a solution of compound **23** (140 mg, 0.33 mmol) in DCM (2 mL) was added TFA (1 mL). The mixture was stirred for 2 h at room temperature. After TLC monitoring, the mixture was concentrated in vacuo and directly used in the next step without further puriﬁcation. The above crude was dissolved in 2 mL DCM, followed by adding Et_3_N (33 mg, 0.33 mmol), tert-butyl 4-formylpiperidine-1-carboxylate (70 mg, 0.33 mmol), and NaBH(OAc)_3_ (140 mg, 0.66 mmol). The mixture was stirred at room temperature overnight. After completion of the reaction, the mixture was quenched with saturated aqueous NaHCO_3_ solution. The mixture was extracted with ethyl acetate. The combined organic layers were washed with brine, dried over Na_2_SO_4_ and concentrated. The crude product was purified by flash column chromatography to afford compound **24** (89 mg, 51 % yield) as a white solid. ^1^H NMR (600 MHz, CDCl_3_) δ 7.95 (s, 1H), 7.73 (d, *J* = 8.6 Hz, 1H), 6.99 (dd, *J* = 8.6, 2.2 Hz, 1H), 6.87 (d, *J* = 2.2 Hz, 1H), 5.20 (dd, *J* = 13.3, 5.1 Hz, 1H), 4.41 (d, *J* = 15.6 Hz, 1H), 4.26 (d, *J* = 15.6 Hz, 1H), 4.10 (s, 3H), 3.36 – 3.28 (m, 4H), 2.94 – 2.78 (m, 2H), 2.71 (s, 1H), 2.57 (t, *J* = 5.0 Hz, 4H), 2.38 – 2.27 (m, 1H), 2.26 – 2.16 (m, 3H), 1.78 – 1.71 (m, 2H), 1.69 – 1.60 (m, 1H), 1.46 (d, *J* = 1.3 Hz, 13H), 1.15 – 1.05 (m, 2H). ESI [M + H]^+^ (*m*/*z*): 526.5.

**tert-Butyl 4-(4-(((S)-1-((2S,4S)-4-hydroxy-2-(((S)-1-(4-(4-methylthiazol-5-yl)phenyl)ethyl)carbamoyl)pyrrolidin-1-yl)-3,3-dimethyl-1-oxobutan-2-yl)amino)-4-oxobutyl)piperidine-1-carboxylate (26)**

To a solution of **25** (50 mg, 0.11 mmol) and 4-(1-(tert-butoxycarbonyl)piperidin-4-yl)butanoic acid (29 mg, 0.11 mmol) in 2 mL DCM were added HATU (46 mg, 0.12 mmol) and DIPEA (71 mg, 0.55 mmol), and the reaction was stirred at room temperature overnight. The reaction mixture was poured into a saturated NH_4_Cl solution, and the aqueous layer was extracted with ethyl acetate. The combined organic layers were washed with brine, dried over Na_2_SO_4_, and concentrated. The crude product was puriﬁed by ﬂash column chromatography to aﬀord the compound **26** (60 mg, 80 % yield) as a white solid.^1^H NMR (600 MHz, CDCl_3_) δ 8.69 (s, 1H), 7.46 (d, *J* = 6.3 Hz, 1H), 7.41 – 7.33 (m, 4H), 6.01 – 5.95 (m, 1H), 5.51 (d, *J* = 9.9 Hz, 1H), 4.71 (d, *J* = 9.1 Hz, 1H), 4.65 (dd, *J* = 14.9, 7.0 Hz, 1H), 4.53 (d, *J* = 9.1 Hz, 1H), 4.47 (dd, *J* = 9.7, 4.8 Hz, 1H), 4.31 (dd, *J* = 14.9, 5.0 Hz, 1H), 4.05 (s, 2H), 3.93 (dd, *J* = 11.0, 4.2 Hz, 1H), 3.82 (dd, *J* = 11.0, 1.6 Hz, 1H), 2.65 (s, 2H), 2.52 (s, 3H), 2.36 (d, *J* = 14.1 Hz, 1H), 2.18 (t, *J* = 7.6 Hz, 3H), 1.84 (d, *J* = 16.4 Hz, 1H), 1.68 – 1.58 (m, 4H), 1.50 – 1.41 (m, 11H), 1.39 – 1.33 (m, 1H), 1.26 – 1.22 (m, 2H), 1.11 – 1.02 (m, 2H), 0.92 (s, 11H). ESI [M + H]^+^ (*m*/*z*): 684.8.

**tert-Butyl (6-((2-(1-methyl-2,6-dioxopiperidin-3-yl)-1,3-dioxoisoindolin-4-yl)amino)hexyl)carbamate (28)**

To a solution of **27** (50 mg, 0.17 mmol) in DMSO (2 mL) was added tert-butyl (6-aminohexyl)carbamate (37 mg, 0.17 mmol) and DIPEA (110 mg, 0.85 mmol). The mixture was stirred at 90 °C overnight. The mixture was diluted with 20 mL of ethyl acetate, and the solution was washed with water (20 mL × 3). The organic layers were washed with brine (20 mL), dried over Na_2_SO_4_, filtered, and concentrated in vacuo. The crude product was puriﬁed by silica gel column chromatography to aﬀord the compound **28** (69 mg, 83 % yield) as a yellow solid. ^1^H NMR (600 MHz, CDCl_3_) δ 7.50 (dd, *J* = 8.5, 7.1 Hz, 1H), 7.10 (d, *J* = 7.1 Hz, 1H), 6.88 (d, *J* = 8.5 Hz, 1H), 6.24 (t, *J* = 5.6 Hz, 1H), 4.96 – 4.90 (m, 1H), 4.55 (s, 1H), 3.27 (td, *J* = 7.1, 5.6 Hz, 2H), 3.22 (s, 3H), 3.16 – 3.10 (m, 2H), 3.03 – 2.93 (m, 1H), 2.83 – 2.72 (m, 2H), 2.15 – 2.06 (m, 1H), 1.68 (p, *J* = 7.2 Hz, 2H), 1.56 – 1.34 (m, 15H). ESI [M + H]^+^ (*m*/*z*): 487.7.

**(2S,4R)-4-Hydroxy-1-((S)-2-(4-(4-((2R,4aS,6aS,12bR,14aS,14bR)-10-hydroxy-2,4a,6a,9,12b,14a-hexamethyl-11-oxo-1,2,3,4,4a,5,6,6a,11,12b,13,14,14a,14b-tetradecahydropicene-2-carbonyl)piperazin-1-yl)butanamido)-3,3-dimethylbutanoyl)-N-((S)-1-(4-(4-methylthiazol-5-yl)phenyl)ethyl)pyrrolidine-2-carboxamide (A1)**

Following the general procedure C, compound **A1** (5 mg, 45%) was obtained as red solid.

^1^H NMR (600 MHz, CDCl_3_) δ 8.67 (s, 1H), 7.53 (d, *J* = 7.8 Hz, 1H), 7.39 (q, *J* = 8.3 Hz, 4H), 7.05 (dd, *J* = 7.1, 1.4 Hz, 1H), 6.93 (s, 1H), 6.59 (d, *J* = 1.4 Hz, 1H), 6.35 (d, *J* = 7.3 Hz, 1H), 5.09 (p, *J* = 7.0 Hz, 1H), 4.74 (t, *J* = 8.1 Hz, 1H), 4.57 (d, *J* = 8.6 Hz, 1H), 4.50 (s, 1H), 4.17 (d, *J* = 11.5 Hz, 1H), 3.74 – 3.55 (m, 4H), 2.53 (s, 4H), 2.44 – 2.27 (m, 9H), 2.22 (s, 3H), 2.14 – 2.08 (m, 2H), 1.92 – 1.65 (m, 10H), 1.56 – 1.43 (m, 12H), 1.25 (s, 6H), 1.12 (s, 3H), 1.05 (s, 9H), 0.99 – 0.93 (m, 1H), 0.60 (s, 3H). ESI [M + H]^+^ (*m*/*z*): 534.26.

**(2S,4R)-4-Hydroxy-1-((S)-2-(6-(4-((2R,4aS,6aS,12bR,14aS,14bR)-10-hydroxy-2,4a,6a,9,12b,14a-hexamethyl-11-oxo-1,2,3,4,4a,5,6,6a,11,12b,13,14,14a,14b-tetradecahydropicene-2-carbonyl)piperazin-1-yl)hexanamido)-3,3-dimethylbutanoyl)-N-((S)-1-(4-(4-methylthiazol-5-yl)phenyl)ethyl)pyrrolidine-2-carboxamide (A2)**

Following the general procedure C, compound **A2** (6 mg, 50%) was obtained as red solid.

^1^H NMR (600 MHz, CDCl_3_) δ 8.67 (s, 1H), 7.50 (d, *J* = 7.9 Hz, 1H), 7.43 – 7.37 (m, 4H), 7.04 (dd, *J* = 7.1, 1.4 Hz, 1H), 6.85 (d, *J* = 8.5 Hz, 1H), 6.52 (d, *J* = 1.5 Hz, 1H), 6.35 (d, *J* = 7.2 Hz, 1H), 5.10 (p, *J* = 7.0 Hz, 1H), 4.75 (t, *J* = 7.9 Hz, 1H), 4.63 (d, *J* = 8.8 Hz, 1H), 4.51 (s, 1H), 4.12 (d, *J* = 11.4 Hz, 1H), 3.74 – 3.54 (m, 1H), 2.57 – 2.51 (m, 4H), 2.47 – 2.31 (m, 9H), 2.21 (s, 3H), 2.15 – 2.03 (m, 2H), 1.88 – 1.56 (m, 12H), 1.54 – 1.44 (m, 10H), 1.39 – 1.30 (m, 4H), 1.26 (d, *J* = 4.9 Hz, 6H), 1.13 (s, 3H), 1.03 (s, 9H), 0.97 (d, *J* = 14.1 Hz, 1H), 0.62 (s, 3H). ESI [M + H]^+^ (*m*/*z*): 534.26.

**(2S,4R)-4-Hydroxy-1-((S)-2-(4-(1-((2R,4aS,6aS,12bR,14aS,14bR)-10-hydroxy-2,4a,6a,9,12b,14a-hexamethyl-11-oxo-1,2,3,4,4a,5,6,6a,11,12b,13,14,14a,14b-tetradecahydropicene-2-carbonyl)piperidin-4-yl)butanamido)-3,3-dimethylbutanoyl)-N-((S)-1-(4-(4-methylthiazol-5-yl)phenyl)ethyl)pyrrolidine-2-carboxamide (A3)**

Following the general procedure C, compound **A3** (7 mg, 47%) was obtained as red solid. ^1^H NMR (600 MHz, CDCl_3_) δ 8.67 (s, 1H), 7.48 – 7.33 (m, 5H), 7.06 (d, *J* = 7.1 Hz, 1H), 6.34 (d, *J* = 7.1 Hz, 1H), 5.09 (p, *J* = 7.0 Hz, 1H), 4.77 (s, 1H), 4.59 (d, *J* = 8.4 Hz, 1H), 4.50 (s, 1H), 4.28 (s, 1H), 3.59 – 3.53 (m, 1H), 3.16 (h, *J* = 3.6 Hz, 1H), 2.53 (s, 3H), 2.35 (t, *J* = 13.4 Hz, 4H), 2.23 – 2.05 (m, 7H), 1.92 – 1.79 (m, 6H), 1.70 – 1.42 (m, 18H), 1.32 – 1.22 (m, 10H), 1.11 (s, 3H), 1.05 (s, 9H), 0.97 – 0.94 (m, 1H), 0.66 (s, 3H). ESI [M + H]^+^ (*m*/*z*): 1030.8.

**(2S,4R)-4-Hydroxy-1-((S)-2-(2-(1-((2R,4aS,6aS,12bR,14aS,14bR)-10-hydroxy-2,4a,6a,9,12b,14a-hexamethyl-11-oxo-1,2,3,4,4a,5,6,6a,11,12b,13,14,14a,14b-tetradecahydropicene-2-carbonyl)azetidin-3-yl)acetamido)-3,3-dimethylbutanoyl)-N-((S)-1-(4-(4-methylthiazol-5-yl)phenyl)ethyl)pyrrolidine-2-carboxamide (A4)**

Following the general procedure C, compound **A4** (4 mg, 33%) was obtained as red solid. ^1^H NMR (600 MHz, CDCl_3_) δ 8.68 (S, 1H), 7.42 – 7.34 (m, 4H), 7.08 (d, *J* = 7.8 Hz, 1H), 6.93 (d, *J* = 27.8 Hz, 1H), 6.57 (d, *J* = 14.8 Hz, 1H), 6.46 (d, *J* = 16.1 Hz, 1H), 6.37 (d, *J* = 7.6 Hz, 1H), 5.10 (p, *J* = 6.9 Hz, 1H), 4.61 – 4.50 (m, 3H), 4.01 (d, *J* = 11.2 Hz, 1H), 3.61 (dd, *J* = 11.3, 3.4 Hz, 1H), 2.82 (d, *J* = 31.7 Hz, 1H), 2.60 – 2.46 (m, 5H), 2.38 – 2.31 (m, 1H), 2.25 – 2.14 (s, 5H), 2.07 – 2.02 (m, 1H), 1.89 – 1.68 (m, 7H), 1.65 – 1.41 (m, 12H), 1.27 – 1.23 (m, 6H), 1.21 – 1.09 (m, 6H), 1.02 (s, 9H), 0.96 (d, *J* = 13.6 Hz, 1H), 0.65 – 0.62 (m, 3H). ESI [M + H]^+^ (*m*/*z*): 974.8.

**(2S,4R)-4-Hydroxy-1-((S)-2-(2-(4-((1-((2R,4aS,6aS,12bR,14aS,14bR)-10-hydroxy-2,4a,6a,9,12b,14a-hexamethyl-11-oxo-1,2,3,4,4a,5,6,6a,11,12b,13,14,14a,14b-tetradecahydropicene-2-carbonyl)piperidin-4-yl)methyl)piperazin-1-yl)acetamido)-3,3-dimethylbutanoyl)-N-((S)-1-(4-(4-methylthiazol-5-yl)phenyl)ethyl)pyrrolidine-2-carboxamide (A5)**

Following the general procedure C, compound **A5** (4 mg, 40%) was obtained as red solid.

^1^H NMR (600 MHz, CDCl_3_) δ 8.67 (s, 1H), 7.88 (d, *J* = 8.2 Hz, 1H), 7.47 (d, *J* = 7.9 Hz, 1H), 7.44 – 7.33 (m, 4H), 7.03 (dd, *J* = 7.1, 1.4 Hz, 1H), 6.96 (s, 1H), 6.50 (s, 1H), 6.35 (d, *J* = 7.2 Hz, 1H), 5.08 (t, *J* = 7.2 Hz, 1H), 4.77 (t, *J* = 7.9 Hz, 1H), 4.50 (d, *J* = 4.1 Hz, 1H), 4.43 (d, *J* = 8.2 Hz, 1H), 4.34 (s, 1H), 4.21 (d, *J* = 11.5 Hz, 1H), 3.58 (dd, *J* = 11.5, 3.6 Hz, 1H), 3.06 – 2.96 (m, 2H), 2.63 – 2.51 (m, 8H), 2.50 – 2.40 (m, 2H), 2.41 – 2.30 (m, 2H), 2.22 – 2.12 (m, 6H), 2.12 – 2.05 (m, 2H), 1.80 – 1.52 (m, 20H), 1.50 – 1.43 (m, 7H), 1.35 – 1.23 (m, 6H), 1.14 (s, 3H), 1.07 (s, 9H), 1.00 – 0.94 (m, 1H), 0.56 (s, 3H). ESI [M + H]^+^ (*m*/*z*): 1100.7.

**(2S,4R)-4-Hydroxy-1-((S)-2-(2-(9-((2R,4aS,6aS,12bR,14aS,14bR)-10-hydroxy-2,4a,6a,9,12b,14a-hexamethyl-11-oxo-1,2,3,4,4a,5,6,6a,11,12b,13,14,14a,14b-tetradecahydropicene-2-carbonyl)-3,9-diazaspiro[5.5]undecan-3-yl)acetamido)-3,3-dimethylbutanoyl)-N-((S)-1-(4-(4-methylthiazol-5-yl)phenyl)ethyl)pyrrolidine-2-carboxamide (A6)**

Following the general procedure C, compound **A6** (5 mg, 42%) was obtained as red solid.

^1^H NMR (600 MHz, CDCl_3_) δ 8.67 (s, 1H), 7.87 (d, *J* = 8.5 Hz, 1H), 7.43 – 7.34 (m, 5H), 7.04 (dd, *J* = 7.2, 1.5 Hz, 1H), 6.95 – 6.90 (m, 1H), 6.50 (d, *J* = 1.4 Hz, 1H), 6.36 (d, *J* = 7.2 Hz, 1H), 5.08 (t, *J* = 7.2 Hz, 1H), 4.75 (t, *J* = 7.9 Hz, 1H), 4.54 – 4.47 (m, 2H), 4.16 (d, *J* = 11.5 Hz, 1H), 3.72 (s, 1H), 3.59 (dd, *J* = 11.4, 3.6 Hz, 1H), 3.02 (s, 2H), 2.57 – 2.46 (s, 8H), 2.35 (dd, *J* = 23.0, 14.6 Hz, 3H), 2.21 (s, 3H), 2.17 (d, *J* = 13.2 Hz, 1H), 2.13 – 2.06 (m, 3H), 1.87 – 1.66 (m, 11H), 1.60 – 1.52 (m, 7H), 1.50 – 1.44 (m, 6H), 1.26 (d, *J* = 4.6 Hz, 6H), 1.14 (s, 3H), 1.07 (s, 9H), 0.97 (d, *J* = 13.9 Hz, 1H), 0.58 (s, 3H). ESI [M + H]^+^ (*m*/*z*): 1071.8.

**(2S,4R)-4-Hydroxy-1-((S)-2-(2-(2-((2R,4aS,6aS,12bR,14aS,14bR)-10-hydroxy-2,4a,6a,9,12b,14a-hexamethyl-11-oxo-1,2,3,4,4a,5,6,6a,11,12b,13,14,14a,14b-tetradecahydropicene-2-carbonyl)-2,7-diazaspiro[3.5]nonan-7-yl)acetamido)-3,3-dimethylbutanoyl)-N-((S)-1-(4-(4-methylthiazol-5-yl)phenyl)ethyl)pyrrolidine-2-carboxamide (A7)**

Following the general procedure C, compound **A7** (6 mg, 44%) was obtained as red solid.

^1^H NMR (600 MHz, CDCl_3_) δ 8.68 (s, 1H), 7.83 (d, *J* = 8.7 Hz, 1H), 7.43 – 7.35 (m, 5H), 7.06 (dd, *J* = 7.2, 1.4 Hz, 1H), 6.91 (s, 1H), 6.48 (d, *J* = 1.4 Hz, 1H), 6.37 (d, *J* = 7.2 Hz, 1H), 5.09 (p, *J* = 7.0 Hz, 1H), 4.75 (t, *J* = 8.0 Hz, 1H), 4.58 – 4.51 (m, 2H), 4.12 – 4.03 (m, 3H), 3.64 – 3.57 (m, 2H), 3.50 – 3.44 (m, 1H), 3.06 – 2.92 (m, 2H), 2.54 – 2.46 (m, 6H), 2.27 (d, *J* = 14.1 Hz, 1H), 2.24 – 2.12 (m, 6H), 2.07 (td, *J* = 14.0, 3.8 Hz, 2H), 1.91 – 1.74 (m, 9H), 1.73 – 1.62 (m, 4H), 1.61 – 1.55 (m, 3H), 1.53 – 1.43 (m, 6H), 1.27 (s, 3H), 1.16 (s, 3H), 1.11 (s, 3H), 1.06 (s, 9H), 1.01 – 0.93 (m, 1H), 0.64 (s, 3H). ESI [M + H]^+^ (*m*/*z*): 1043.8.

**(2R,4aS,6aS,12bR,14aS,14bR)-N-(4-((2-(2,6-Dioxopiperidin-3-yl)-1,3-dioxoisoindolin-4-yl)amino)butyl)-10-hydroxy-2,4a,6a,9,12b,14a-hexamethyl-11-oxo-1,2,3,4,4a,5,6,6a,11,12b,13,14,14a,14b-tetradecahydropicene-2-carboxamide (A8)**

Following the general procedure C, compound **A8** (10 mg, 55%) was obtained as red solid.

^1^H NMR (600 MHz, CDCl_3_) δ 8.26 (s, 1H), 7.46 (t, *J* = 7.7 Hz, 1H), 7.08 (d, *J* = 6.9 Hz, 1H), 7.00 (d, *J* = 7.0 Hz, 1H), 6.84 (d, *J* = 8.4 Hz, 1H), 6.53 (s, 1H), 6.33 (d, *J* = 6.8 Hz, 1H), 6.19 (s, 1H), 5.80 (s, 1H), 4.94 – 4.89 (m, 1H), 3.28 – 3.09 (m, 4H), 2.92 – 2.69 (m, 3H), 2.45 (d, *J* = 15.4 Hz, 1H), 2.20 (s, 3H), 2.12 (d, *J* = 6.6 Hz, 2H), 2.02 (td, *J* = 14.2, 3.7 Hz, 1H), 1.96 – 1.82 (m, 4H), 1.70 – 1.41 (m, 16H), 1.14 (s, 3H), 1.12 (s, 3H), 1.02 (dt, *J* = 14.1, 3.6 Hz, 1H), 0.63 (s, 3H). ESI [M + H]^+^ (*m*/*z*): 777.5.

**(2R,4aS,6aS,12bR,14aS,14bR)-N-(6-((2-(2,6-Dioxopiperidin-3-yl)-1,3-dioxoisoindolin-4-yl)amino)hexyl)-10-hydroxy-2,4a,6a,9,12b,14a-hexamethyl-11-oxo-1,2,3,4,4a,5,6,6a,11,12b,13,14,14a,14b-tetradecahydropicene-2-carboxamide (A9)**

Following the general procedure C, compound **A9** (8 mg, 52%) was obtained as red solid.

^1^H NMR (600 MHz, CDCl_3_) δ 8.34 (d, *J* = 49.7 Hz, 1H), 7.50 – 7.45 (m, 1H), 7.08 (d, *J* = 7.0 Hz, 1H), 7.00 (d, *J* = 7.1 Hz, 1H), 6.85 (d, *J* = 8.5 Hz, 1H), 6.53 (s, 1H), 6.34 (d, *J* = 7.1 Hz, 1H), 6.19 (d, *J* = 5.0 Hz, 1H), 5.70 (t, *J* = 3.0 Hz, 1H), 4.91 (ddd, *J* = 12.3, 5.3, 1.6 Hz, 1H), 3.21 (q, *J* = 6.7 Hz, 2H), 3.16 – 3.04 (m, 2H), 2.93 – 2.68 (m, 3H), 2.43 (dt, *J* = 15.1, 2.8 Hz, 1H), 2.21 – 2.14 (m, 4H), 2.03 (td, *J* = 14.3, 3.9 Hz, 1H), 1.97 – 1.81 (m, 4H), 1.71 – 1.29 (m, 21H), 1.14 (s, 3H), 1.12 (s, 3H), 1.02 (dd, *J* = 13.9, 3.6 Hz, 1H), 0.64 (d, *J* = 1.4 Hz, 3H). ESI [M + H]^+^ (*m*/*z*): 805.2.

**(2R,4aS,6aS,12bR,14aS,14bR)-N-(8-((2-(2,6-Dioxopiperidin-3-yl)-1,3-dioxoisoindolin-4-yl)amino)octyl)-10-hydroxy-2,4a,6a,9,12b,14a-hexamethyl-11-oxo-1,2,3,4,4a,5,6,6a,11,12b,13,14,14a,14b-tetradecahydropicene-2-carboxamide (A10)**

Following the general procedure C, compound **A10** (9 mg, 50%) was obtained as red solid.

^1^H NMR (600 MHz, CDCl_3_) δ 8.38 (s, 1H), 7.48 (dd, *J* = 8.5, 7.1 Hz, 1H), 7.08 (d, *J* = 7.1 Hz, 1H), 7.01 (dt, *J* = 7.1, 1.5 Hz, 1H), 6.86 (d, *J* = 9.0 Hz, 1H), 6.54 (t, *J* = 1.5 Hz, 1H), 6.34 (d, *J* = 7.2 Hz, 1H), 6.21 (s, 1H), 5.68 (q, *J* = 5.3 Hz, 1H), 4.92 (m, 1H), 3.22 (t, *J* = 6.9 Hz, 2H), 3.16 – 3.03 (m, 2H), 2.92 – 2.84 (m, 1H), 2.84 – 2.69 (m, 2H), 2.46 – 2.37 (m, 1H), 2.20 (d, *J* = 2.3 Hz, 3H), 2.17 – 2.10 (m, 2H), 2.07 – 1.98 (m, 1H), 1.97 – 1.82 (m, 4H), 1.71 – 1.33 (m, 18H), 1.30 – 1.26 (m, 6H), 1.14 (s, 3H), 1.12 (s, 3H), 1.01 (m, 1H), 0.63 (s, 3H). ESI [M + H]^+^ (*m*/*z*): 833.1.

**(2R,4aS,6aS,12bR,14aS,14bR)-N-(10-((2-(2,6-Dioxopiperidin-3-yl)-1,3-dioxoisoindolin-4-yl)amino)decyl)-10-hydroxy-2,4a,6a,9,12b,14a-hexamethyl-11-oxo-1,2,3,4,4a,5,6,6a,11,12b,13,14,14a,14b-tetradecahydropicene-2-carboxamide (A11)**

Following the general procedure C, compound **A11** (8 mg, 43%) was obtained as red solid.

^1^H NMR (600 MHz, CDCl_3_) δ 8.41 (d, *J* = 45.6 Hz, 1H), 7.53 – 7.47 (m, 1H), 7.10 (d, *J* = 7.0 Hz, 1H), 7.02 (d, *J* = 7.1 Hz, 1H), 6.89 (d, *J* = 8.5 Hz, 1H), 6.55 (d, *J* = 3.9 Hz, 1H), 6.35 (d, *J* = 7.1 Hz, 1H), 6.24 (s, 1H), 4.94 (dd, *J* = 12.3, 5.2 Hz, 1H), 3.29 – 3.05 (m, 3H), 2.93 – 2.72 (m, 3H), 2.22 (d, *J* = 2.2 Hz, 3H), 2.15 (dt, *J* = 15.0, 3.9 Hz, 2H), 2.05 (td, *J* = 14.3, 3.6 Hz, 1H), 1.99 – 1.84 (m, 3H), 1.72 – 1.30 (m, 18H), 1.16 (s, 3H), 1.14 (s, 3H), 0.65 (s, 3H). ESI [M + H]^+^ (*m*/*z*): 861.6.

**2-(2,6-Dioxopiperidin-3-yl)-4-((1-((2R,4aS,6aS,12bR,14aS,14bR)-10-hydroxy-2,4a,6a,9,12b,14a-hexamethyl-11-oxo-1,2,3,4,4a,5,6,6a,11,12b,13,14,14a,14b-tetradecahydropicene-2-carbonyl)piperidin-4-yl)ethynyl)isoindoline-1,3-dione (A12)**

Following the general procedure C, compound **A12** (3 mg, 29%) was obtained as red solid. ^1^H NMR (600 MHz, CDCl_3_) δ 8.07 (s, 1H), 7.87 – 7.62 (m, 3H), 7.05 – 6.90 (m, 2H), 6.47 (s, 1H), 6.36 (d, *J* = 7.2 Hz, 1H), 5.00 – 4.96 (m, 1H), 3.06 – 2.69 (m, 4H), 2.38 – 2.31 (m, 3H), 2.21 (s, 3H), 2.20 – 2.03 (m, 4H), 1.88 – 1.59 (m, 15H), 1.45 (s, 3H), 1.30 (d, *J* = 1.1 Hz, 6H), 1.16 (s, 3H), 1.04 – 0.97 (m, 2H), 0.63 (s, 3H). ESI [M + H]^+^ (*m*/*z*): 798.6.

**2-(2,6-dioxopiperidin-3-yl)-4-((2-(1-((2R,4aS,6aS,12bR,14aS,14bR)-10-hydroxy-2,4a,6a,9,12b,14a-hexamethyl-11-oxo-1,2,3,4,4a,5,6,6a,11,12b,13,14,14a,14b-tetradecahydropicene-2-carbonyl)piperidin-4-yl)ethyl)amino)isoindoline-1,3-dione (A13)**

Following the general procedure C, compound **A13** (4 mg, 38%) was obtained as red solid.

^1^H NMR (600 MHz, CDCl_3_) δ 7.50 (dd, *J* = 8.4, 7.1 Hz, 1H), 7.13 – 7.09 (m, 1H), 7.04 (m, 1H), 6.86 (dd, *J* = 8.5, 3.0 Hz, 1H), 6.54 (d, *J* = 14.8 Hz, 1H), 6.35 (dd, *J* = 7.3, 2.5 Hz, 1H), 6.20 (dt, *J* = 24.0, 5.4 Hz, 1H), 4.95 – 4.92 (m, 1H), 4.31 (d, *J* = 13.0 Hz, 1H), 3.32 (m, 2H), 2.97 – 2.69 (m, 4H), 2.37 (dd, *J* = 39.4, 14.2 Hz, 2H), 2.25 – 2.07 (m, 6H), 1.87 – 1.49 (m, 21H), 1.45 (s, 3H), 1.30 – 1.26 (m, 5H), 1.13 (d, *J* = 4.3 Hz, 3H), 0.98 (d, *J* = 14.2 Hz, 1H), 0.50 (s, 3H). ESI [M + H]^+^ (*m*/*z*): 817.5.

**2-(2,6-Dioxopiperidin-3-yl)-4-((4-(1-((2R,4aS,6aS,12bR,14aS,14bR)-10-hydroxy-2,4a,6a,9,12b,14a-hexamethyl-11-oxo-1,2,3,4,4a,5,6,6a,11,12b,13,14,14a,14b-tetradecahydropicene-2-carbonyl)piperidin-4-yl)butyl)amino)isoindoline-1,3-dione (A14)**

Following the general procedure C, compound **A14** (3 mg, 30%) was obtained as red solid.

^1^H NMR (600 MHz, CDCl_3_) δ 7.50 – 7.47 (m, 1H), 7.09 (dd, *J* = 7.0, 1.7 Hz, 1H), 7.05 – 7.01 (m, 1H), 6.88 (d, *J* = 8.4 Hz, 1H), 6.52 (d, *J* = 4.9 Hz, 1H), 6.35 (d, *J* = 7.2 Hz, 1H), 6.29 – 6.22 (m, 1H), 4.94 – 4.90 (m, 1H), 4.31 (t, *J* = 7.0 Hz, 1H), 3.28 (m, 2H), 2.94 – 2.70 (m, 4H), 2.36 (dd, *J* = 34.0, 13.3 Hz, 2H), 2.22 (d, *J* = 8.4 Hz, 3H), 2.18 – 2.07 (m, 3H), 1.89 – 1.48 (m, 21H), 1.45 – 1.41 (m, 5H), 1.29 – 1.25 (m, 7H), 1.13 (d, *J* = 2.2 Hz, 3H), 0.97 (d, *J* = 14.1 Hz, 1H), 0.53 (s, 3H). ESI [M + H]^+^ (*m*/*z*): 845.6.

**2-(2,6-Dioxopiperidin-3-yl)-4-(4-((1-((2R,4aS,6aS,12bR,14aS,14bR)-10-hydroxy-2,4a,6a,9,12b,14a-hexamethyl-11-oxo-1,2,3,4,4a,5,6,6a,11,12b,13,14,14a,14b-tetradecahydropicene-2-carbonyl)piperidin-4-yl)methyl)piperazin-1-yl)isoindoline-1,3-dione (A15)**

Following the general procedure C, compound **A15** (6 mg, 43%) was obtained as red solid.

^1^H NMR (600 MHz, CDCl_3_) δ 8.26 (d, *J* = 56.9 Hz, 1H), 7.59 (t, *J* = 7.7 Hz, 1H), 7.39 (d, *J* = 7.1 Hz, 1H), 7.17 (d, *J* = 8.3 Hz, 1H), 7.03 (d, *J* = 7.1 Hz, 1H), 6.53 (s, 1H), 6.35 (d, *J* = 7.2 Hz, 1H), 4.96 (ddd, *J* = 12.6, 5.5, 1.5 Hz, 1H), 4.35 (s, 1H), 3.36 (d, *J* = 12.7 Hz, 4H), 2.94 – 2.58 (m, 8H), 2.43 – 2.07 (m, 12H), 1.91 – 1.65 (m, 11H), 1.61 – 1.49 (m, 4H), 1.46 (s, 3H), 1.33 (d, *J* = 14.4 Hz, 1H), 1.29 (s, 3H), 1.27 (s, 3H), 1.14 (s, 3H), 0.98 (dt, *J* = 14.0, 3.6 Hz, 1H), 0.57 (s, 3H). ESI [M + H]^+^ (*m*/*z*): 872.6.

**2-(2,6-Dioxopiperidin-3-yl)-5-(4-((1-((2R,4aS,6aS,12bR,14aS,14bR)-10-hydroxy-2,4a,6a,9,12b,14a-hexamethyl-11-oxo-1,2,3,4,4a,5,6,6a,11,12b,13,14,14a,14b-tetradecahydropicene-2-carbonyl)piperidin-4-yl)methyl)piperazin-1-yl)isoindoline-1,3-dione (A16)**

Following the general procedure C, compound **A16** (5 mg, 39%) was obtained as red solid.

^1^H NMR (600 MHz, CDCl_3_) δ 8.16 (s, 1H), 7.69 (d, *J* = 8.4 Hz, 1H), 7.29 (d, *J* = 2.2 Hz, 1H), 7.05 (ddd, *J* = 14.6, 7.8, 1.8 Hz, 2H), 6.52 (s, 1H), 6.36 (d, *J* = 7.2 Hz, 1H), 4.94 (dd, *J* = 12.5, 5.4 Hz, 1H), 4.36 (s, 1H), δ 3.41 (t, *J* = 5.1 Hz, 4H), 2.91 – 2.67 (m, 4H), 2.57 (dq, *J* = 11.5, 5.6 Hz, 4H), 2.42 – 2.06 (m, 12H), 1.92 – 1.66 (m, 11H), 1.61 – 1.49 (m, 4H), 1.46 (s, 3H), 1.36 – 1.31 (m, 1H), 1.29 (s, 3H), 1.27 (s, 3H), 1.14 (s, 3H), 0.58 (s, 3H). ESI [M + H]^+^ (*m*/*z*): 858.6.

**3-(5-(3-(4-((2R,4aS,6aS,12bR,14aS,14bR)-10-Hydroxy-2,4a,6a,9,12b,14a-hexamethyl-11-oxo-1,2,3,4,4a,5,6,6a,11,12b,13,14,14a,14b-tetradecahydropicene-2-carbonyl)piperazin-1-yl)prop-1-yn-1-yl)-1-oxoisoindolin-2-yl)piperidine-2,6-dione (A17)**

Following the general procedure C, compound **A17** (4 mg, 35%) was obtained as yellow solid. ^1^H NMR (600 MHz, CDCl_3_) δ 8.02 (s, 1H), 7.81 (t, *J* = 7.5 Hz, 1H), 7.52 (q, *J* = 6.9 Hz, 2H), 7.01 (dd, *J* = 7.0, 1.4 Hz, 2H), 6.48 (dd, *J* = 8.2, 1.4 Hz, 1H), 6.35 (d, *J* = 7.2 Hz, 1H), 5.23 – 5.19 (m, 1H), 4.49 – 4.46 (m, 1H), 4.35 – 4.31 (m, 1H), 3.53 (s, 2H), 2.97 – 2.79 (m, 2H), 2.60 (s, 4H), 2.40 – 2.32 (m, 4H), 2.26 – 2.05 (m, 7H), 1.88 – 1.64 (m, 11H), 1.44 (s, 3H), 1.30 – 1.29 (m, 6H), 1.15 (s, 3H), 1.01 – 0.94 (m, 2H), 0.60 (d, *J* = 6.1 Hz, 3H). ESI [M + H]^+^ (*m*/*z*): 799.5

**3-(5-(4-((1-((2R,4aS,6aS,12bR,14aS,14bR)-10-Hydroxy-2,4a,6a,9,12b,14a-hexamethyl-11-oxo-1,2,3,4,4a,5,6,6a,11,12b,13,14,14a,14b-tetradecahydropicene-2-carbonyl)piperidin-4-yl)methyl)piperazin-1-yl)-1-oxoisoindolin-2-yl)piperidine-2,6-dione (A18)**

Following the general procedure C, compound **A18** (5 mg, 45%) was obtained as yellow solid.

^1^H NMR (600 MHz, CDCl_3_) δ 8.07 (s, 1H), 7.73 (d, *J* = 8.6 Hz, 1H), 7.04 (d, *J* = 6.9 Hz, 1H), 6.99 (d, *J* = 8.6 Hz, 1H), 6.88 (s, 1H), 6.52 (s, 1H), 6.36 (d, *J* = 7.1 Hz, 1H), 5.20 (dd, *J* = 13.2, 5.0 Hz, 1H), 4.41 (d, *J* = 15.6 Hz, 1H), 4.26 (d, *J* = 15.6 Hz, 1H), 3.31 (t, *J* = 5.2 Hz, 4H), 2.92 – 2.77 (m, 3H), 2.61 – 2.54 (m, 4H), 2.42 – 2.08 (m, 12H), 1.91 – 1.69 (m, 12H), 1.62 – 1.49 (m, 4H), 1.46 (s, 3H), 1.27 (t, *J* = 10.2 Hz, 7H), 1.14 (s, 3H), 1.01 – 0.94 (m, 1H), 0.57 (s, 3H). ESI [M + H]^+^ (*m*/*z*): 872.6.

**(2S,4S)-4-Hydroxy-1-((S)-2-(4-(1-((2R,4aS,6aS,12bR,14aS,14bR)-10-hydroxy-2,4a,6a,9,12b,14a-hexamethyl-11-oxo-1,2,3,4,4a,5,6,6a,11,12b,13,14,14a,14b-tetradecahydropicene-2-carbonyl)piperidin-4-yl)butanamido)-3,3-dimethylbutanoyl)-N-((S)-1-(4-(4-methylthiazol-5-yl)phenyl)ethyl)pyrrolidine-2-carboxamide (A3-NC)**

Following the general procedure C, compound **A3-NC** (8 mg, 47%) was obtained as red solid. ^1^H NMR (600 MHz, CDCl_3_) δ 8.68 (s, 1H), 7.38 (q, *J* = 8.2 Hz, 5H), 7.06 – 7.02 (m, 1H), 6.52 (s, 1H), 6.34 (d, *J* = 7.2 Hz, 1H), 5.53 (d, *J* = 9.9 Hz, 1H), 4.78 (s, 1H), 4.66 (dd, *J* = 14.9, 7.1 Hz, 1H), 4.55 (d, *J* = 8.8 Hz, 1H), 4.49 – 4.46 (m, 1H), 4.32 (dd, *J* = 14.9, 5.0 Hz, 1H), 3.94 (dd, *J* = 11.1, 4.2 Hz, 1H), 3.81 (dd, *J* = 11.0, 1.6 Hz, 1H), 2.52 (s, 3H), 2.41 – 2.28 (m, 4H), 2.21 – 2.16 (m, 7H), 1.91 – 1.41 (m, 24H), 1.34 – 1.21 (m, 10H), 1.12 (s, 3H), 0.98 – 0.94 (m, 2H), 0.90 (s, 9H), 0.66 (s, 3H). ESI [M + H]^+^ (*m*/*z*): 1016.9.

**(2R,4aS,6aS,12bR,14aS,14bR)-10-Hydroxy-2,4a,6a,9,12b,14a-hexamethyl-N-(6-((2-(1-methyl-2,6-dioxopiperidin-3-yl)-1,3-dioxoisoindolin-4-yl)amino)hexyl)-11-oxo-1,2,3,4,4a,5,6,6a,11,12b,13,14,14a,14b-tetradecahydropicene-2-carboxamide (A9-NC)**

Following the general procedure C, compound **A9-NC** (8 mg, 52%) was obtained as red solid. ^1^H NMR (600 MHz, CDCl_3_) δ 7.48 (dd, *J* = 8.5, 7.1 Hz, 1H), 7.08 (d, *J* = 7.0 Hz, 1H), 7.01 – 6.95 (m, 2H), 6.85 (d, *J* = 8.5 Hz, 1H), 6.52 (s, 1H), 6.33 (d, *J* = 7.1 Hz, 1H), 6.19 (t, *J* = 5.6 Hz, 1H), 5.69 – 5.67 (m, 1H), 4.94 – 4.88 (m, 1H), 3.22 – 3.18 (m, 5H), 3.17 – 3.03 (m, 2H), 3.02 – 2.93 (m, 1H), 2.79 – 2.70 (m, 2H), 2.44 (d, *J* = 15.5 Hz, 1H), 2.19 (s, 3H), 2.15 – 2.07 (m, 2H), 2.03 – 2.00 (m, 1H), 1.97 – 1.82 (m, 4H), 1.68 – 1.30 (m, 20H), 1.14 (s, 3H), 1.12 (s, 3H), 1.04 – 0.99 (m, 1H), 0.64 (s, 3H). ESI [M + H]^+^ (*m*/*z*): 534.26. ESI [M + H]^+^ (*m*/*z*): 819.8.
